# Contribution of the neuron-specific ATP1A3 to embryonic spinal circuit emergence

**DOI:** 10.1101/2024.06.11.598491

**Authors:** Sarah Dinvaut, Sophie Calvet, Jean-Christophe Comte, Raphael Gury, Olivier Pascual, Maelys André, Rosaria Ferrigno, Jérôme Honnorat, Frédéric Moret, Guillaume Marcy, Julien Falk, Valérie Castellani

**Author notes:** Co-first authors (Sarah Dinvaut, Sophie Calvet). Co-last authors (Julien Falk, Valérie Castellani). Corresponding authors emails S. Dinvaut V. Castellani J. Falk.

## Abstract

The early neurodevelopmental contributions of ion pumps remain poorly characterized. Combining analysis of public human embryo single-cell transcriptomic datasets and an embryonic chicken model, we found a conserved differentiation sequence whereby spinal cord neurons switch on neuron-specific alpha3 subunit (ATP1A3) of Na^+^/K^+^ ATPases. In the chicken model, ATP1A3 is distributed along axons and growth cones. Its knockdown alters axon pathfinding of dorsal interneurons (DIN) that wire spinocerebellar circuits. In mirror of reported electric field (EF)-driven cell migration, we found that DIN axons align in EFs, which was abolished by Na^+^/K^+^ ATPase inhibitor Ouabain and ATP1A3 knockdown. We recorded an embryonic trans-neural-epithelial potential generating EF whose pharmacological and surgical manipulation mimicked ATP1A3 knock-down-induced altered DIN axon pathfinding. Using DINs transplantation paradigm, we found that ATP1A3 is required cell-autonomously for EF-mediated long-range guidance. Finally, dominant-negative ATP1A3 mutation causing an early ATP1A3 childhood disease disrupts this fundamental developmental process, revealing unexpected pathogenic mechanisms.

## Introduction

Ion balance and fluxes across the plasma membrane are regulated by ions channels and pumps. While channels let ions flow along their electrochemical gradient, pumps can actively transport ions against their gradient and are therefore essential to regulate ion balance and channel activity. Although ion channel activity is generally considered to be the preserve of mature excitable cells, these molecules are also important for immature and non-excitable cells, through yet largely unclear mechanisms. During development, ion channel expressions are detected in various tissues and their loss of function can have dramatic effects ^1,2^. Mutations in ion channels cause several human syndromes characterized by craniofacial and limb development alterations. Furthermore, ion channel modulations by antiepileptic drugs during gestation leads to congenital malformations ^2–5^. In the developing nervous system, progenitors and immature neurons express channels that usually differ from the ones found in mature cells. Some early contributions were reported, ranging from neural tissue patterning, progenitor division, migration to neuron differentiation and cell death, both in the brain and spinal cord ^6–11^. Studies from different animal models also support a contribution of ion channels to axon elongation and pathfinding ^12–17^. These key steps of circuit formation ensure that the developing axons connect their appropriate target.

Among ionic pumps, Na^+^/K^+^ ATPases that use ATP to export 3Na^+^ and import 2 K^+^, are major regulators of the ionic balance and cell resting potential. Na^+^/K^+^ ATPases pumps are composed of a catalytic alpha (α) subunit, a beta (β) regulatory subunit and eventually another accessory regulatory subunit FXYD. Four genes encode alpha subunits (*ATP1A1-4*) that differ in expression, Na^+^ affinity and Ouabain sensitivity ^18–21^.

Seminal works in Xenopus showed that tight regulation of Na^+^/K^+^ ATPases activity, through β3 subunit, is required for neuronal specification during the early stages of neural tube formation^22,23^. In Zebrafish larvae, alpha subunits were shown to be indispensable for brain ventricle expansion ^24,25^. In human, the neuronal specific ATP1A3 gene is expressed in the fetal cortex and mutations can cause neurological diseases that are manifested shortly after birth ^21,26–32^. Although these diseases were commonly viewed as primarily due to aberrant excitability of postnatal neurons, ATP1A3 mutations can lead to cortical malformations probably resulting from impaired migration of cortical neuron in fetuses ^27,28,30^.

Here, we thought to characterize ATP1A3 roles at early stages of embryonic spinal cord neural circuit formation. Through analyses of single-cell RNA sequencing (scRNAseq) dataset in human and of expression patterns in chick, we report that neuronal differentiation is marked by activation of ATP1A3 gene expression. In the chicken embryo experimental model, we show that its loss of function impairs the stereotypic navigation of dorsal interneurons (DINs) that establishes circuits with brain structures. Next, we investigated how Na^+^/K^+^ ATPases could control axons guidance. We postulated that they could be involved in a mechanism of guidance by electric fields (EF), based on literature reporting that Electric field (EF) provide positional information ^33–37^ during embryogenesis and that Na^+^/K^+^ ATPases mediate EF-directed migration of cells ^38^. Using a range of *in vitro* and *in vivo* approaches we found that ATP1A3 controls axon response to embryonic electric field (EF) generated by a transneural-epithelial potential. Finally, we found that overexpression of the mutated form of ATP1A3 (E815K) causing the most severe forms of the pediatric neurological disease called alternative hemiplegia of childhood (AHC)^32^ alters axon navigation of DINs. This suggests that ATP1A3 dysfunctions during early steps of spinal cord-brain circuit formation could contribute to AHC etiology.

## Results

### ATP1A3 is expressed in early postmitotic neurons of the human spinal cord

To gain first insights into ATP1A3 early contributions, we studied its expression profile in the human embryonic spinal cord. Different neurons subtypes emerge from specific progenitor pools that organize along the dorso-ventral axis of the spinal cord (Figure 1a). Previous works showed that similarly to other species, human dorsal interneurons are generated after motoneurons and appear after the Carnegie stages CS13 (gestational weeks {GW} 4.5) but are reliably detectable at stage CS18 (GW7)^39,40^. We analyzed the single-cell RNAseq datasets generated by Rayon and collaborators^41^ at CS14 (GW5) and CS17 (GW6) using standard unsupervised clustering that was recently shown to efficiently identify spinal neuron and progenitor subpopulations from single-cell data of human embryonic spinal cord^42^. Cells were categorized into several progenitor and neuronal clusters and identified with well-recognized markers at both stages (Sup figure 1a-h). Consistently with the analysis of Rayon and collaborators^41^, we found that neuron proportion rose from 41% to 63% between CS14 (n=2,909 cells) and CS17 (n=6,004) (Sup figure 1b), with increase by more than 2-fold of dorsal interneurons and specification of additional sub-types (Sup figure 1d and e). We therefore focused on CS17 for the study of ATP1A3 expression. We identified progenitor and neuron clusters (Figure 1a and b, Sup figure 1f) and conducted a differential gene expression analysis. As expected, genes related to proliferation are down regulated in neurons while those related to neuronal differentiation are upregulated (Figure 1c). Interestingly, we found ATP1A3 among genes whose expression is significantly upregulated in neurons (Log2(FC)=1.15, adjusted p-value=1.66*10^-169^) (Figure1 c). ATP1A3 expression was detected in all neuronal clusters (NEFM positive) (figure1 d), including the Dl1 neuron cluster identified at CS17 (Sup figure 1g and h). ATP1A3 expression rose by more than 2-fold in DI1 neurons compared to the Dl1-2 progenitors (log2(FC)=1.015, adjusted p-value= 4.668*10^-14^) (Sup figure 1h). In conclusion, ATP1A3 is expressed in early differentiating neurons of the human spinal cord.

**Figure 1:**
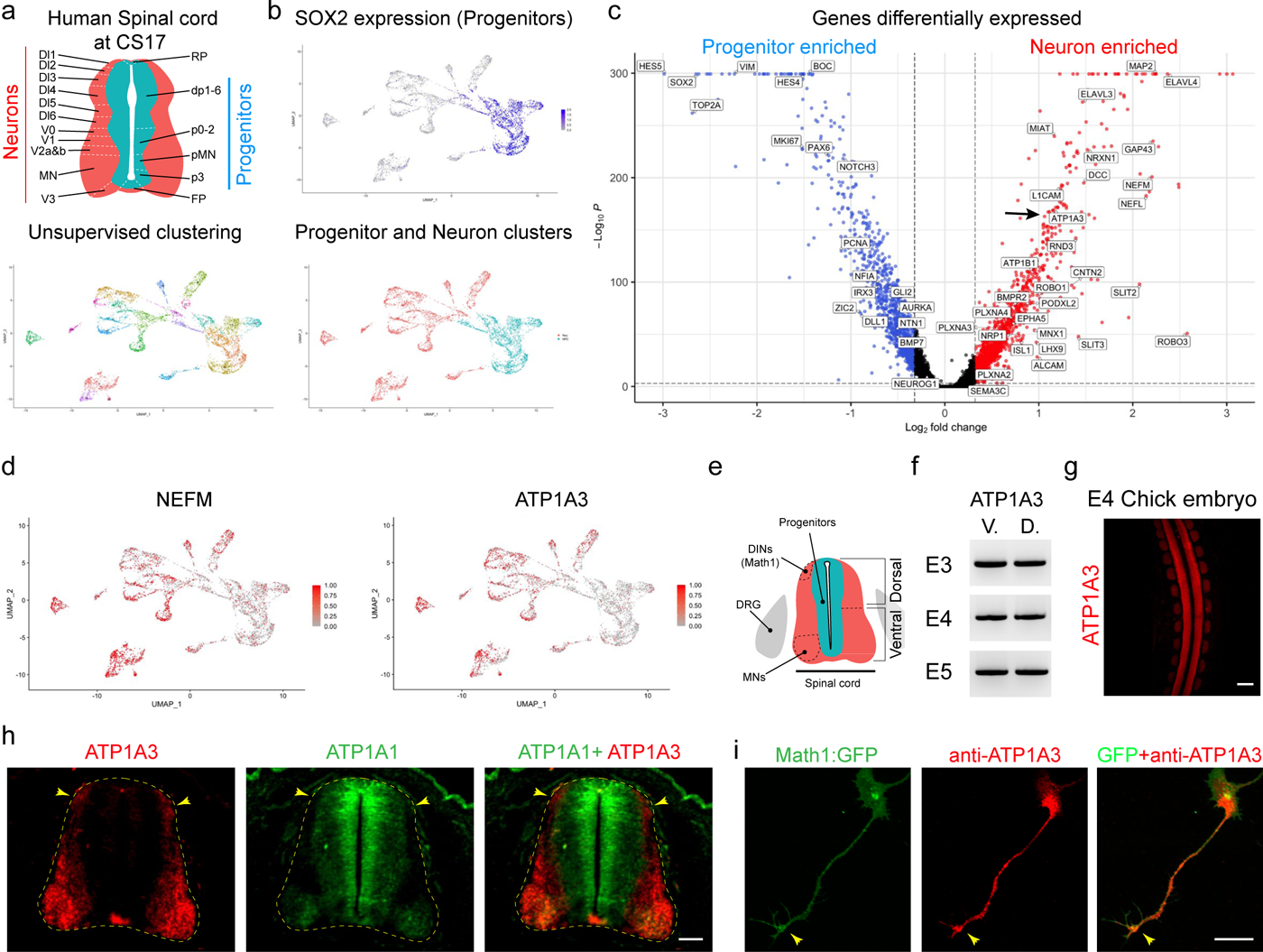
Human and chicken embryonic spinal cord neurons expressed ATP1A3 during their morphological differentiation. (a) scheme of neuron and progenitor subtypes domain in the human spinal cord at CS17 (top) and UMAP of different clusters identified (bottom). b) Feature plot showing the expression of SOX2 that identifies progenitor populations (top). UMAP of neuronal and progenitor clusters at CS17 (bottom). (c) Volcano plot of the genes differentially expressed between progenitors and neurons. Genes with higher expression in neurons are shown in red and in blue when lower (i.e. higher in progenitors). The names of several genes relevant for progenitor status, neuronal differentiation and guidance cue receptors are indicated. Arrow points to ATP1A3. (d) Feature plot showing the cell expressing the medium neurofilament (NEFM) (left) and ATP1A3 (right). (e) Scheme of the chicken embryonic spinal cord depicting the different regions and position of motoneurons (MN) and dorsal interneurons (DINs). (f) RT-PCR detection of ATP1A3 transcripts in dorsal (D.) and ventral (V.) spinal cord at E4, E5 and E6. (g) Top view of the ATP1A3 expression detected by HCR RNA-FISH in whole mount E4 embryo. (h) Virtual coronal section of E4 spinal cord embryo showing in red ATP1A3 expressions in MN and DINs (arrowheads) and in green ATP1A1 expression in progenitors and MNs. The dashed line outlines the spinal cord. (i) Immunolabelling of ATP1A3 on primary dorsal spinal cord culture. ATP1A3 (red) is found on the axon and growth cone (arrowhead) of DINs expressing GFP (green) under the DI1-specific promoter Math1. Scale bars: 300µm (g), 50µm (h), 20µm (i).

### ATP1A3 is expressed in chicken spinal neurons as they form their axons

To study ATP1A3 functions in the spinal cord, we took advantage of the chicken embryo model that is amenable to various experimental manipulations. CS17, the stage at which we found ATP1A3 expression activated in spinal cord neurons is considered to correspond to mouse E11.5 ^41^ or chicken E4^43^ developmental stages marked by axon navigation of spinal neurons. This suggested that ATP1A3 could contribute to neuronal differentiation and axon development.

To assess the conservation of ATP1A3 expression, we investigated its expression profile in the spinal cord of chicken embryo (Figure 1e). We could confirm using RT-PCR that ATP1A3 subunit is expressed in both ventral (MN) and dorsal (DIN) spinal cord from E3 to E5 (Figure 1f). To determine if ATP1A3 is selectively expressed by differentiating neurons, we performed HRC RNA-FISH on whole mount chicken embryos. We found that ATP1A3 is expressed in the E4 spinal cord and that, in contrast to ATP1A1, its expression is restricted to the mantle layer, where post-mitotic neurons differentiate (Figure 1g-h). Thus, similarly to human spinal cord, ATP1A3 appears to be switched on in chick embryonic spinal neurons. We also observed that in contrast to motoneurons (MNs), dorsal interneurons (DINs) do not express ATP1A1 (Figure 1h). We focused on these neurons that may be particularly sensitive to ATP1A3 perturbation as they do not express ATP1A1.

We performed immunostaining of α3 in primary cultures of dorsal spinal cord tissue dissected out from embryos electroporated with Math1::GFP plasmid that restricts GFP expression to DINs. We detected α3 subunit in the axons and the growth cones of DIN neurons (Figure 1i). Altogether, the embryonic DINs represented a promising population to assess the contributions of ATP1A3 to early processes of neuronal differentiation.

### ATP1A3 expression contributes to the navigation of DIN axons

To investigate the role of ATP1A3 in DINs, we knocked-down ATP1A3 gene expression by electroporating a custom siRNA in E2 chicken embryos. SiRNA efficacy was first validated in neurons dissociated from E3 spinal cord. By quantifying the intensity of α3 immunolabeling in electroporated cells identified by GFP expression, we found that SiRNA at a concentration 1.25µg/µl reduced α3 expression by 76% (Sup figure 2a and b, Control SiRNA: 88038±13233, N=21 cells; ATP1A3 siRNA: 20756±4441, N=16 cells; p<0.0001). After 24h *in vitro*, differentiating neurons transfected with the ATP1A3 siRNA exhibited reduced α3 expression in their processes compared to the control siRNA condition (Sup figure 2c).

**Figure 2:**
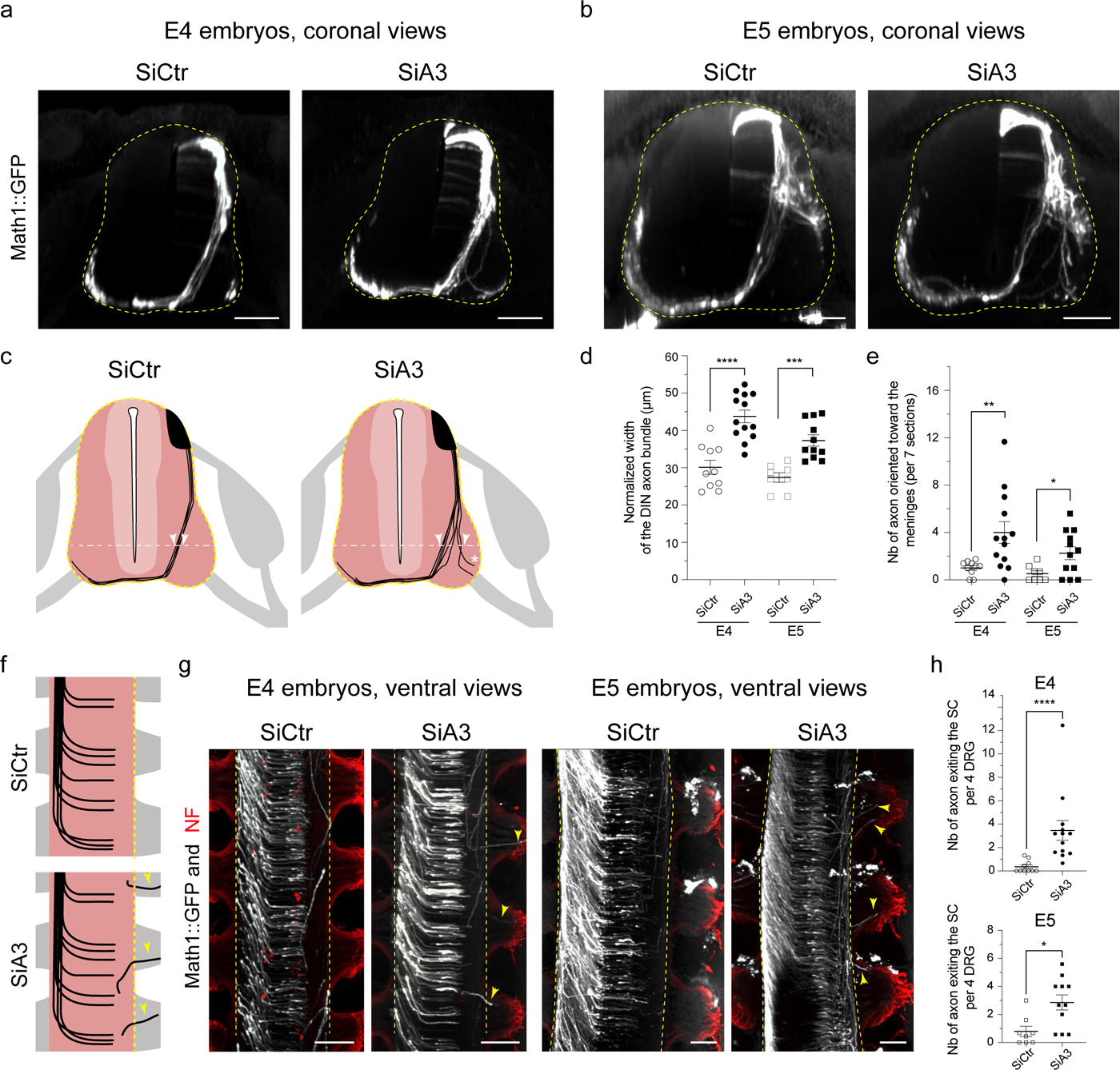
ATP1A3 knock-down impair DIN axon navigation to the floorplate and their confinement to the spinal cord. (a-b) Coronal views of embryo co-electroporated with Math1::GFP and control (SiCtr) or siRNA targeting ATP1A3 (SiA3) at E4 (a) and E5 (b). Yellow dashed lines outline the spinal cord. (c) Drawings schematizing in black the trajectories observed for DIN axons in SiCtr and SiA3 embryos on coronal views of the spinal cord. White dashed lines mark the level 1/5 of spinal cord width used for quantification in (d). White arrows highlight medial- and lateral-most positions of DIN axons. (d-e) Scatter dot plots showing the DIN axon bundle width (d) and the number of axons abnormally orientating toward the meninges (e) in SiCtr and SiA3 in E4 and E5 embryos. (f) Drawings illustrating the different types of trajectories observed in longitudinal view for DINs axons transfected with SiCtr (top) and SiA3 (bottom). (g) Longitudinal views of the ventral spinal cord after co-electroporation of SiA3 or SiCtr with Math1::GFP in E4 and E5 embryos. (h) Scatter dot plots showing the number of axons exiting the spinal cord in each condition. Bars indicate the mean ± SEM, *p<0.05, **p < 0.01; (Mann-Whitney). S.C Spinal cord. GFP staining is shown in white and neurofilament (NF) in red (g). Scale bars: 100µm (a and b) 150µm (g).

To analyze the contribution of ATP1A3 to the development of DIN axons, we electroporated the siRNAs with Math1::GFP plasmid that specifically drives GFP expression in DIN neurons. 48h and 72h after electroporation, the embryos were cleared to analyze DIN axon trajectories with light sheet microscopy. This technique enables to fully appreciate 3D axon organization and to reveal phenotypes that would be difficult to identify otherwise ^44,45^. We found after control or ATP1A3 targeting siRNA electroporation that GFP+ neurons formed and extended projections that have a pattern characteristic of the DIN population with axons extending ventrally to cross the midline in the floor plate (Figure 2a-c; Sup figure 2d and e). However, at both stage (E4 and E5), after ATP1A3 knock-down GFP+ axons exhibited abnormal trajectories. In the control siRNA condition, DIN axons properly shifted from a lateral to more medial position when reaching the ventral part of the spinal cord (Figure 2a-c, left panels). In contrast, in the ATP1A3-siRNA condition, DIN axons often diverted from this stereotypical trajectory. Many failed to switch to a normal medial position with some orientating toward the meninges (Figure 2a-c, right panels). This resulted in an increase of the mediolateral dispersion of DIN axons (Figure 2d; E4 SiCtr: 30.11 ± 1.86 µm, N=10, E4 SiA3: 43.76 ± 1.69 µm, N=13, p<0.0001; E5 SiCtr: 27.38 ± 1.25 µm, N=8, E5 SiA3= 37.30 ± 1.53 µm, N=11, p=0.0001, Mann Whitney) and a rise of axons aberrantly turning toward the meninges (Figure 2e; E4 SiCtr: 1.01 ± 0.190, N=10, E4 SiA3=4.00 ± 0.91, N=13, p=0.0048; E5 SiCtr: 0.54±0.23, N=8, E5 SiA3: 2.258±0.55, N=12, p=0.040, Mann Whitney). Strikingly, some axons even exited the spinal cord through ventral roots (Figure 2f-h; E4 SiCtr: 0.36 ± 0.16, N=10, E4 SiA3: 3.46 ± 0.85, N=10, p<0.0001; E5 SiCtr: 0.80 ± 0.37, N=8, E5 SiA3: 2.86 ± 0.54, N=11, p= 0.014, Mann Whitney) (Sup figure 2f). Thus, altering ATP1A3 expression impacted trajectories of DIN axons suggesting that ATP1A3 provides information to properly orient DIN axon navigation and confinement within the spinal cord.

### ATP1A3 mediates EF-induced orientation of DIN axons

Next, we investigated how ATP1A3 could contribute to axon navigation. Na^+^/K^+^ ATPases support EFs-directed migration of cells^38^ and axons from various neurons of different animal species were shown to orient in EFs, *in vitro* ^46–48^. Thus, we hypothesized that ATP1A3 knock-down induced aberrant navigation of DIN axons could result from the loss of guidance information normally provided by EFs.

First, we examined using an *in vitro* set-up whether cultured neurons from dissociated dorsal spinal cord (DIN) orient their axons according to EF. 35, 80 and 140mV/mm EFs were chosen to cover the range of representative endogenous EFs reported in the embryo ^49–54^. To prevent contamination by electrochemical reaction products, platinum electrodes were bathed in 30ml-bottles filled with medium and connected to the culture chamber with agar-salt bridges ^55–58^ (Figure 3a). EFs were applied 1h after dissociated neuron plating for 24h, a set-up that enables the axons to form and elongate. While DIN axons showed random trajectories in absence of EF, they oriented towards the cathode (sink of current, negative pole) when exposed to EF, as reflected by an increase in the orientation index (0 for random orientation to 1 when all axons orient in the same direction; Figure 3b-d). DIN axons responded to EF in a voltage gradient strength-dependent manner (Figure 3e; DIN 0mV: 0.044±0.011, N=9; DIN 35mV: 0.299±0.107, N=3, p=0.1045; DIN 80mV: 0.290±0.030, N=3; p=0.0045; DIN 140mV: 0.450±0.054, N=3; p=0.0045, Mann-Whitney). EFs were shown to regulate various processes, from early cell polarization to neurite initiation and axon turning ^48,59,60^. To test for direct effect on axon growth orientation, we applied EFs 16h after plating, at a time when neurons already extended their axons (Figure 3f). After 8 hours application of 140mV/mm, we observed a significant orientation effect on DIN axons (Figure 3g; DIN 0mV:0.028±0.008, N=4; DIN140mV: 0.299±0.038, N=7; p=0.0061, Mann-Whitney). Thus, EF can control the orientation of DIN axons as they develop.

**Figure 3:**
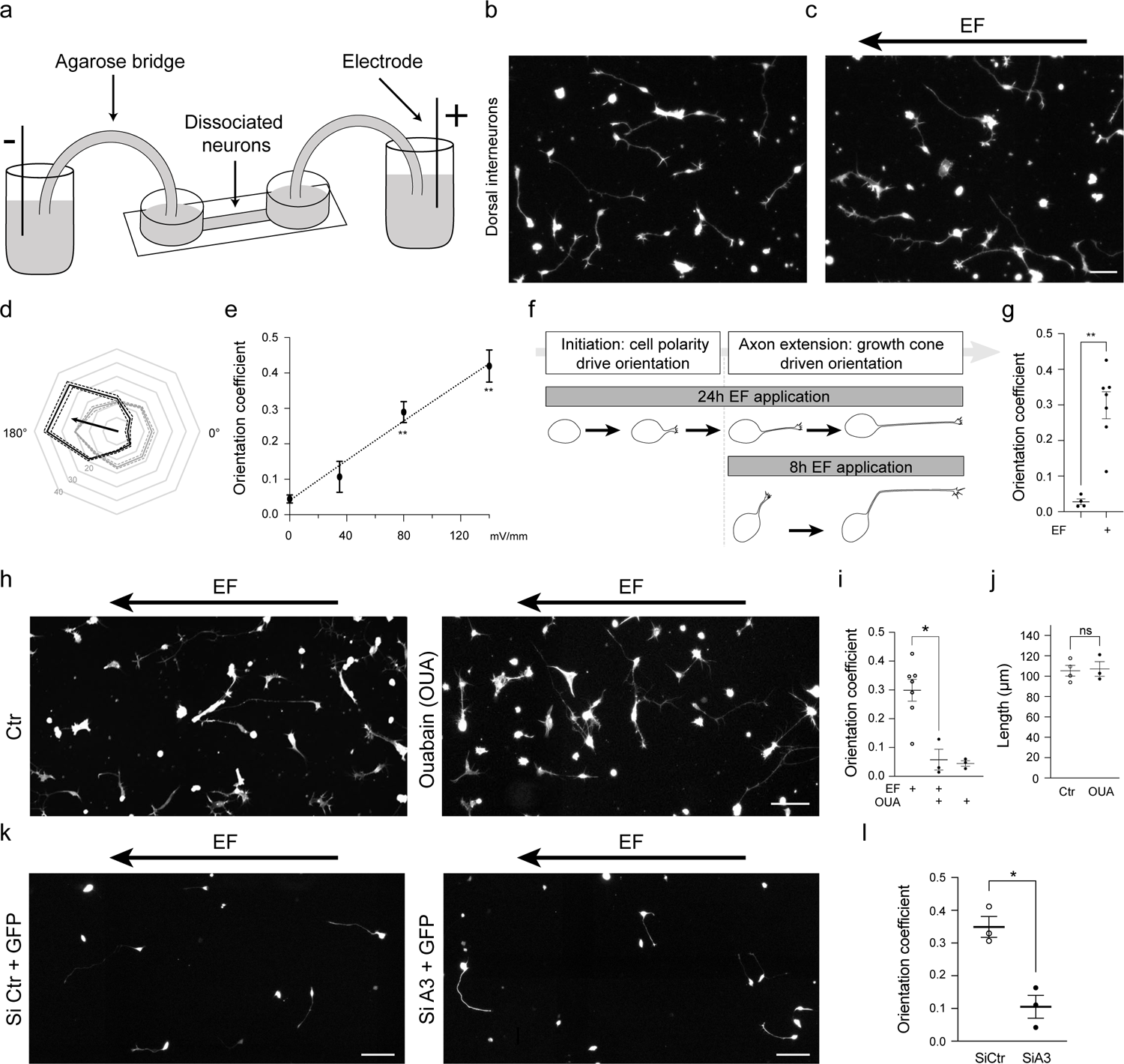
DIN axon orientation in response to Electric fields depends on ATP1A3. (a) Diagram representing the set-up of electric stimulation in cultures of DINs dissociated neurons. (b-c) Representative pictures showing the cathodal (-) tropism of axons after 24h exposure to 140mV electric field (EF) (c) in contrast to control condition (b). Dissociated DINs were labeled with phalloidin (white). (d) Radar plot of the angle distribution for DIN axons exposed to 0mV (grey) or 140mV EF (black). Plain lines indicate the mean values and the dashed lines the SEM values. The black arrow represents the mean vector orientation for DIN axons. (e) Dose response curves of DIN axons to EFs showing the average of the orientation coefficients obtained for independent cultures exposed to 0mV (N=9 DIN), 35mV (N=3 DIN), 80mV (N=3 DIN) and 140mV (N=3 DIN) for 24h. The orientation coefficient is the norm of the mean vector of the axon populations (several hundred of axons per cultures). The norm increases from 0 (random angular distribution of axons) to 1 (same angle for all axons). (f) Drawing illustrating the different phases of axon formation potentially influenced depending on the timings of EF stimulation after plating. In contrast to 24h stimulations, 8h stimulations starting 16h after plating impact axonal extension and not axon initiation. (g) Scatter dot plot showing the orientation coefficients of DIN axons exposed or not to 140mV EF during 8h, 16h after plating. (h) Mosaic images of phalloidin-labeled cultured neurons exposed to EF in absence (Ctr) (left panel) or in presence of Ouabain (OUA) (right panel). (i) Scatter dot plot of the orientation coefficient obtained for cultures exposed to EF in presence or absence of Ouabain (OUA). (j) Scatter dot plot showing the average axon length of neurons cultured in absence (Ctr) or presence of ouabain (OUA). (k) Representative field of GFP+neurons (white) after co-transfection with control SiRNA (siCtr) (left panel) or SiRNA targeting ATP1A3 (SiA3) (right panel). (l) Scatter dot plot showing the orientation coefficient of dissociated neurons transfected with control SiRNA (SiCtr) or SiRNA targeting ATP1A3 (SiA3). Bars indicate the mean ± SEM. *p < 0.05; **p < 0.01; ***p < 0.001 (Mann-Whitney). Scale bars: 50µm(c), 100µm (h and k).

Second, we studied whether Na^+^/K^+^ ATPases are involved in this EF-induced axon orientation. We started by blocking their function with the cardiac glycoside Ouabain (OUA). We observed that application of 10μM Ouabain resulted in increase of intracellular Na^+^ in DIN neurons (Sup figure 2g and h) and 80% decrease of the EF-mediated orientation of their axons (Figure 3h and I; 140mV/mm: 0.29 ± 0.04, N=7; 140mV/mm + OUA (10μM): 0.06 ± 0.04, N=3, p = 0.033; Mann-Whitney). Ouabain application did not impair basal axon growth supporting a specific role of Na^+^/K^+^ ATPases in EF-induced axon trajectories (Figure 3j). Next, we quantified the response of siRNA transfected neurons and found that ATP1A3 siRNA decreased by 71% the response of DINs to EF while control siRNA had no effect (Figure 3k and l; control siRNA: 0.35 ± 0.03, N=3 independent experiments; ATP1A3: 0.10 ± 0.03, N=3 independent experiments; p=0.05, Mann-Whitney). Thus, DINs perceive EFs and orient their axons accordingly, and the mechanism requires α3 containing Na^+^/K^+^ ATPases.

### Extracellular EFs are present in the spinal cord wall and could contribute to DIN axon navigation

Next, we searched for EFs in the chicken embryo that could provide positional information during DIN axon navigation in the spinal cord, with interest for trans-neural tube potential (TNTP), previously characterized in other species ^50,52,54^. First, we measured the TNTP in the tectum of E4 chicken embryos, a region accessible to such experimentation. We recorded the electric potential as we moved progressively the tungsten electrodes across the tectum wall from its surface to ventricle (Figure 4a). We found that the cerebrospinal fluid potential was negative compared to the surface of the tectum (Figure 4b). The electric potential drop was not due to extraembryonic membranes as their removal had no impact (Ctr: 5.8 ± 0.6 mV, N= 29 measures from 10 Embryos; W/O ExM: 9.3 ± 2.0 mV, N= 16 measures from 7 embryos) (Figure 4c). Second to interfere with the TNTP, we thought to perform surgical incision at the hindbrain level to open the neural tube and allow ionic homogenization between the CSF and the extra-embryonic space (Figure 4a, OB). We observed with electrophysiological measurement that it resulted in an 80 % drop of the TNTP (OB: 1.8 ± 0.2 mV; N=42 measures from 16 Embryos; W/O ExM versus OB, p<0.0001, Mann-Whitney) (Figure 4c).

**Figure 4:**
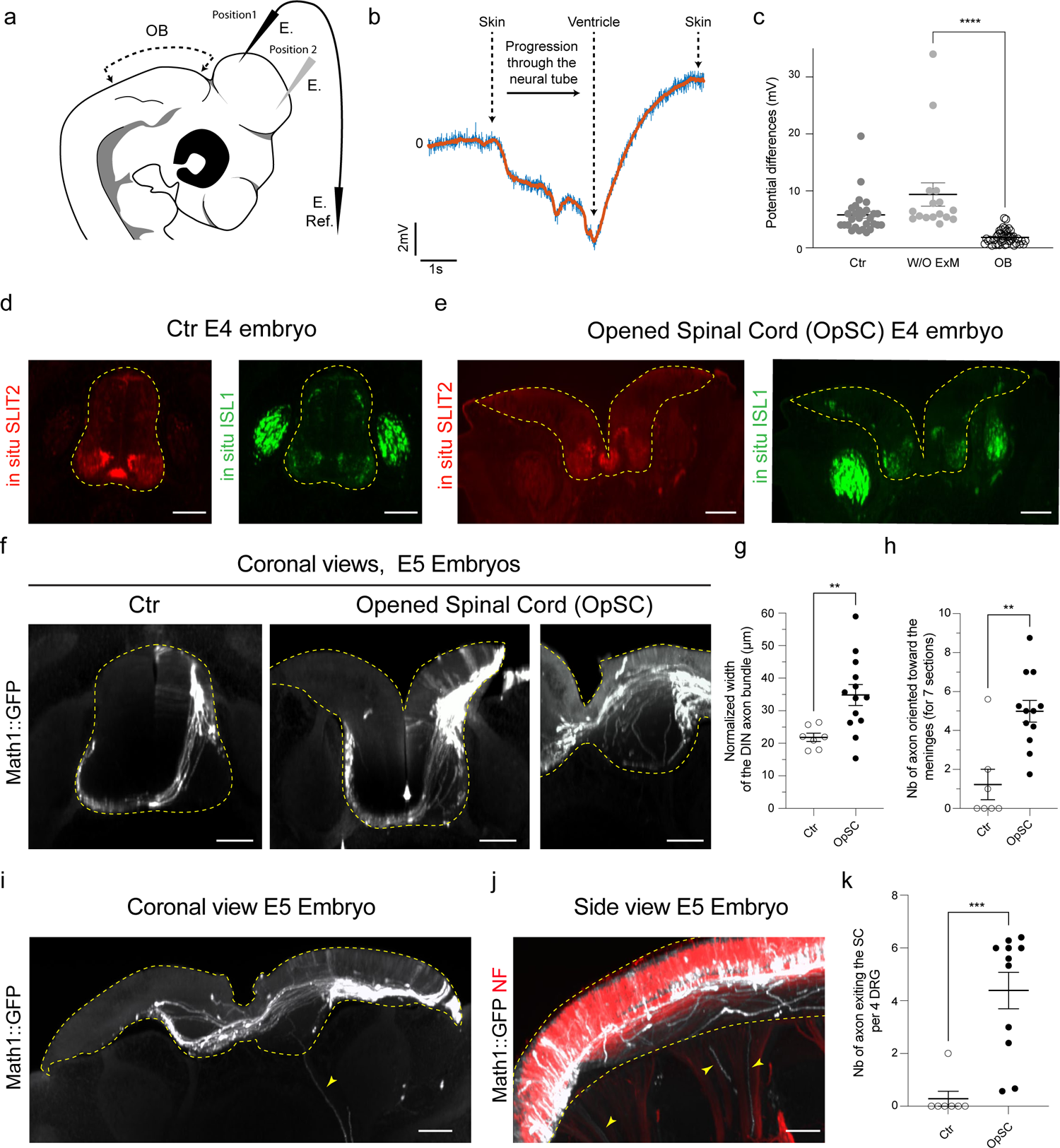
Developing spinal cord exhibits endogenous electric field that could control DIN axon navigation. (a) Drawing illustrating E4 embryo head morphology, recording positions (Electrode, E.; reference electrode E. Ref.) and the surgically opened region (Dashed line, Open brain OB). (b) Example of electrical record of the trans-neural tube potential in the tectum of E4 embryos. (c) Scatter dot plot of the electric potential differences between the surface of the tectum and the ventricle measured in embryos with intact extra-embryonic membranes (Ctr, dark gray circles), without extra-embryonic membrane (W/OExM, grey circles) and having surgical opening at the hindbrain level (OB, open circles). (d-e) Coronal views of HCR in situ fluorescence for SLIT2 and ISL1 (Islet1) in E4 control embryos (d) and in E4 embryos after surgical opening of the spinal cord at E3 (e). (f) Spinal cord coronal views of E5 embryos electroporated with Math1::GFP in control condition (left panel) or opened spinal cord (right panels) showing the changes in medio-lateral positions and orientation of DIN axons. (g-h) Scatter dot plots of the normalized width of DIN axon bundle (g) and of the number of axons oriented toward the meninges (h). (i-j) Coronal view (i) and side view (j) of E5 opened spinal cord embryo showing GFP positive DIN axons exiting the spinal cord (arrowheads). Neurofilament (NF) counterstaining was overlaid in j to highlight the ventral roots. (k) Scatter dot plot of the number of axons exiting the spinal cord normalized for 4 DRG length. Bars indicate the mean ± SEM. **p < 0.01; ***p < 0.001 (Mann-Whitney). Dashed lines in d, e, f, i, j outline the spinal cord. Scale bars: 100µm (d, e, f, i and j).

Third, we assessed the consequences of TNTP reduction resulting from neural tube opening on DIN axon navigation. Thus, we performed surgical opening of the lumbo-thoracic spinal cord on Math1::GFP electroporated embryos at E3. To make sure that spinal cord patterning was maintained and that floor plate functions were preserved, we performed HCR RNA-FISH on E4 operated embryos. Consistently with the fact that *ex vivo* spinal cord open-books can be used to study DIN axon navigation across the floor plate^61,62^, we showed that the surgical manipulation did not interfere with expression of key guidance molecules such as Slit2 or neuron specification such as Isl1 expression (Figure 4d-e). As expected, Isl1 expression was still detected in a dorsal population of neurons (Figure 4d-e). GFP expression was maintained in Math1::GFP transfected neurons. These neurons appeared correctly localized and formed axons (Figure 4 f-h). However, we found that GFP positive axons often deviated from their normal path. Quantifications showed that they have abnormal mediolateral positions in the ventral spinal cord (Figure 4 f-h; Ctr: 21.83 ± 1.23 µm, N=7 embryos; OpSC: 34.87 ± 3.23µm, N=13, p= 0.0063; Meninge oriented Ctr: 1.23 ± 0.78, N=7 embryos; OpSC: 4.99 ± 0.56, N=12, p= 0.0046, Mann Whitney), with some axons also exiting the ventral spinal cord, exactly as they do so after ATP1A3 knock-down (Figure 4i-k; Ctr: 0.28±0.28, N=7 embryos; OpSC: 4.39± 0.69, N=11, p= 0.0002, Mann Whitney). This supports a contribution of EF to DIN axon navigation in the spinal cord.

### ATP1A3 is required cell autonomously for EF-mediated DIN axon navigation

Several mechanisms could underlie ATP1A3 function during DIN axon navigation. Its expression by spinal neurons could be required for setting the TNTP and/or for orienting their axons in the EF. Deciphering ATP1A3 mode of action thus required a paradigm to uncouple the formation of the TNTP and the response to the EF. Interestingly, the TNTP was reported to act over large distances, with crucial contribution to head patterning ^50,54^. Thus, to distinguish between these two potential mechanisms of action, we developed an *in vivo* paradigm consisting in ectopically engrafting DIN neuronal explants outside the spinal cord of host embryos. Axons extending from the explants were thus given the choice to navigate in the three dimensions of the embryo. This paradigm allowed us to assess if they could be specifically oriented toward the spinal cord by the TNTP and if so if this guidance is dependent on their ATP1A3 expression. We performed *in ovo* neural tube electroporation at E2 to express the GFP under the control of DIN-specific enhancer (Math1). One day later we grafted dorsal (DIN) spinal cord explants of donor electroporated embryos, at the base of the lower limb of E3 recipient embryos (Figure 5a). We analyzed the trajectories of GFP^+^ axons two days post-grafting using light-sheet microscopy of transparentized embryos. Remarkably, we observed that DIN axons preferentially oriented towards their natural navigation territory, the spinal cord (Figure 5b). In addition to DIN axons emerging from the transplant with appropriate orientation, we observed axons initially growing in erroneous direction but secondly correcting their trajectory (Figure 5b and Sup Figure 3a). We could also observe GFP+ axons entering and extending in the spinal cord (Sup Figure 3b and c). Notably, after reaching the SC, a vast majority of individual DIN axons turned longitudinally, ascending rostrally towards the brain along a ventral path, as endogenous DIN axons normally do (Sup Figure 3d and e). Such proper axon orientation of transplanted neurons suggested maintenance of their identity in this ectopic environment. In support, we found by immunolabeling that Math1-driven GFP^+^ DIN neurons expressed typical markers of their lineage such as Robo3 and that other markers such as Isl1 was maintained in the graft (Sup Figure 3f-h). We designed qualitative and quantitative methods to evaluate the spatial orientation of axon trajectories at global and individual levels. First, we quantified the orientation of the ten longest DIN axons and found that a large majority extended towards or in the SC (64.5% of axons, n=220 axons, N=21 DIN grafts). Second, we set-up a global orientation index (adapted from Castellani et al., 2000 and Falk et al.,2005 ^63,64^) with a score of 0 for grafts having all axons orienting towards the spinal cord and of 1 when all axons oriented towards the limb. We found a mean orientation index of 0.29±0.03 (N=22 grafts) reflecting their preferential orientation towards their spinal cord (p<0.0001, Mann-Whitney two tail). To assess whether this orientation towards the SC was specific to DIN, we engrafted explants of motoneurons whose projections are drastically different since they project their axons away from the spinal cord towards the periphery. Interestingly, unlike DIN axons, motoneuron axons oriented toward the limb (0.74±0.04 for the N=39 grafts), very rarely entering the spinal cord (Sup Figure 3k-m). Engraftment of motoneuron explants in a second ectopic site had similar outcomes, as the axons still preferentially oriented towards the periphery.

**Figure 5:**
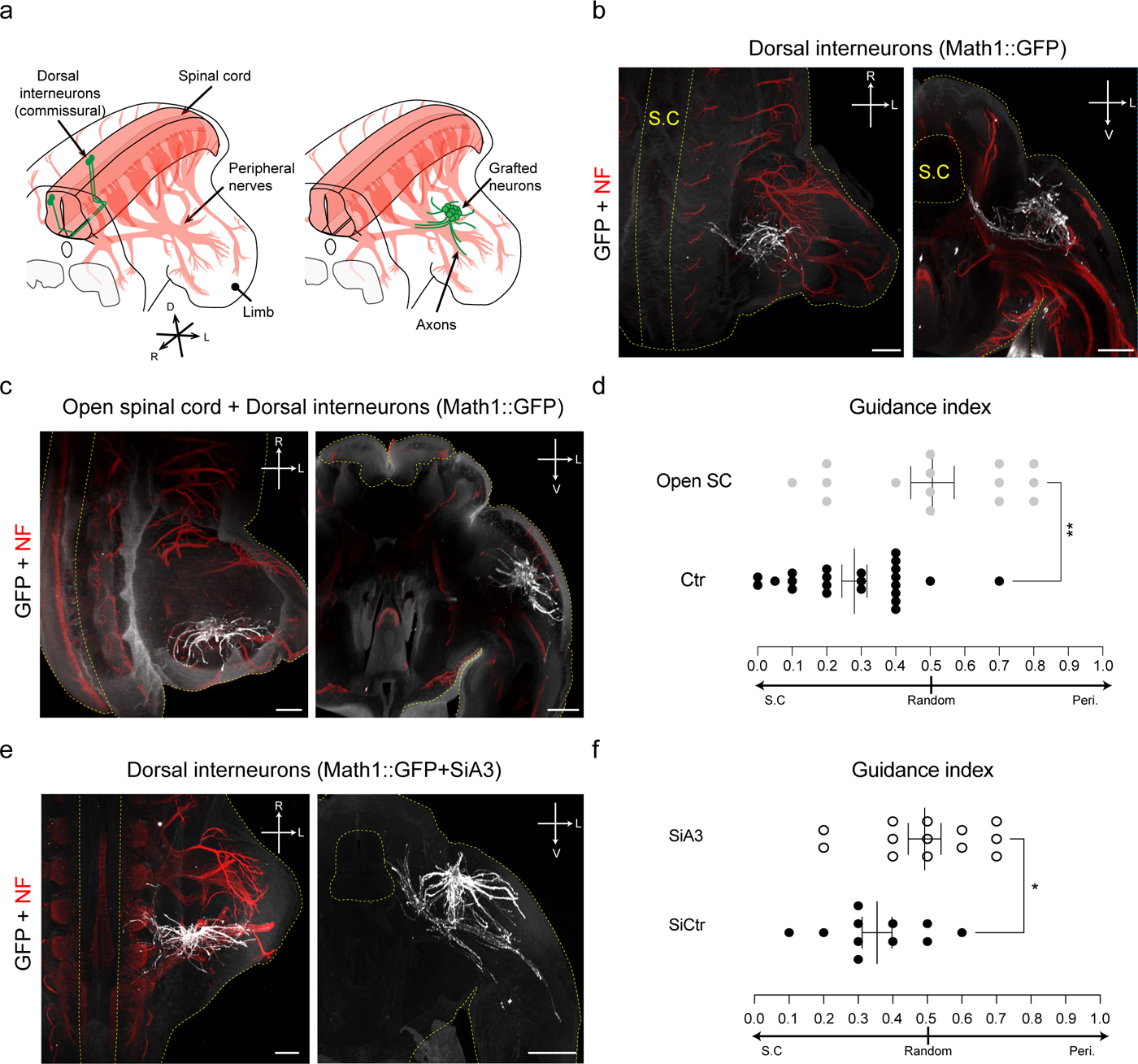
ATP1A3-mediated response to electric fields could contribute to long-range guidance of DIN axon navigating from ectopic position. (a) Diagrams showing the endogenous projections of DIN within the spinal cord of the chick embryo (left) and the grafting paradigm (right). (b-c) 3D top (left) and coronal (right) views exemplifying DIN (Math1::GFP) axon orientation toward the spinal cord (S.C) when neurons are grafted in control embryos (b) in contrast to the random DIN axon orientation observed after spinal cord opening (c). (d) Scatter dot plots showing the guidance index of the axons formed by DINs grafted in embryos without opened spinal cord (Ctr, black circles) or with opened spinal cord (Open SC, grey circles). (e) Top (left panel) and coronal (right) views of an embryo grafted with explant co-electroporated with Math1::GFP and siRNA targeting ATP1A3 (SiA3). (f) Scatter dot plot showing the guidance index for DINs transfected with control siRNA (SiCtr, black) or siRNA targeting ATP1A3 (SiA3, open circles). For all images, GFP staining is shown in white and neurofilament (NF) in red. Yellow dashed lines outline the embryo and the spinal cord (S.C). R, rostral; L, lateral, V ventral. S.C Spinal cord, Peri. Periphery. *p<0.05, **p < 0.01 (Mann-Whitney). Scale bars: 300µm (b, c and e).

Next using this paradigm, we assessed if the orientation of DIN axons towards the spinal cord depends on the TNTP and on ATP1A3 expression in transplanted DIN neurons.

First, we combined spinal cord surgery to abrogate the TNTP and grafting of DIN explants. Two days later, we observed that the neural tube opening was still present and we quantified the orientation of DIN axons towards the spinal cord. We found that DIN axon trajectories were no longer oriented towards the spinal cord, which was reflected by a guidance index that was significantly different from that of our grafts in intact embryos (Figure 5c and d; 0.51±0.06, N=15 grafts; p=0.0031, Mann-Whitney). Thus, neural tube opening both decreased the TNTP and altered DIN axon trajectories. Second, we performed siRNA-mediated knockdown of ATP1A3 in donor embryos prior to transplantation of DINs explants in host embryos. Similarly, it resulted in the loss of spinal cord-oriented navigation observed in the control siRNA condition (Figure 5e and f; control siRNA: 0.35 ± 0.04, N=11 grafts; ATP1A3: 0.49 ± 0.05, N=13 grafts; p=0.0486, Mann-Whitney). Thus, these findings suggest that ATP1A3 is required cell-autonomously for DIN neurons to perceive the TNTP and orient accordingly towards the spinal cord.

### Expression of the ATP1A3 bearing the E815K mutation causing AHC, impairs DIN axon navigation

Lastly, we assessed whether human mutations in ATP1A3 gene could interfere with proper navigation of DIN axons. We focused on E815K mutation, the second most common of ATP1A3 that causes the most severe form of Alternating Hemiplegia of Childhood (AHC) ^32^. We cloned the chicken wild-type ATP1A3 (WT) from E4 spinal cord and introduced a point mutation to reproduce the human E815K substitution. To facilitate identification of transfected cells, tomato was expressed from the same plasmid. First, we studied the impact of the E815K substitution on ionic transport efficiency. In cultures of electroporated dorsal spinal cord cells, we measured intracellular Na+ and K+ using CoroNa green and PSBI fluorescent probes^65,66^, respectively. We found that ATP1A3-WT expression increased Na^+^/K^+^ transport, thus validating the cloning of a functional ATP1A3 (Figure 6a and b; CoroNa Green Tomato: 281.4± 22.97, N=22 fields; ATP1A-WT: 195.1± 21.41, N=20 fields, p= 0.013) (Sup figure 4a and b). Oppositely, ATP1A3-E815K expression decreased Na^+^/K^+^ transport, as expected from the reported dominant negative effect of this substitution^67,68^ (Figure 6c and d ; CoroNa Green Tomato: 182.6 ± 20.91, N=24 fields; ATP1A-815: 326.7 ± 36.63, N=24 fields, p= 0.0008)(Sup Figure 4a and c). Next, we analyzed axon navigation of DIN neurons after co-electroporation of ATP1A3-WT and ATP1A3-E815K in combination with Math1::GFP (Sup figure 4d). Similarly to what we observed with siRNA-mediated knock-down, we found that expression of ATP1A3-E815K altered the mediolateral positioning of DIN axons increasing the spreading along this axis by 1.4 fold (Figure 6e and f). Some axons diverged from the main tract to orient toward the meninges and eventually exiting the spinal cord by ventral roots (Figure 6g-I and Sup figure 4e). Gain of ATP1A3-WT also altered the mediolateral positioning of DIN axons. As a result, the normalized distance between the most medial and lateral axons increased for both ATP1A3 constructs (Figure 6e and f; Tomato: 22.95± 1.12, N=6; ATP1A3-815: 31.92± 1.91, N=13, p= 0.0002; Tomato: 19.20± 1.07, N=10, ATP1A3-WT: 31.13± 1.807, N=15 p= 0.0002, Mann Whitney). However, in contrast to the ATP1A3-E815K condition, axons overexpressing ATP1A3-WT rarely oriented toward the meninges or exited the spinal cord (Figure 6g-j; Meninge-oriented axons, Tomato: 0.44 ± 0.19, N=6; ATP1A3-815: 2.55 ± 0.39, N=13 embryos, p=0.0006; Tomato: 0.64 ± 0,20, N=10, ATP1A3-WT: 1.36 ± 0.38, N=15, p= 0.1755; Exit of the spinal cord, Tomato: 0,2619± 0,1176, N=6; ATP1A3-815: 1.11± 0.26, N=13 embryos, p= 0.041; Tomato: 0.24± 0.13, N=10, ATP1A3-WT: 0.40± 0.15, N=15 p= 0.476, Mann Whitney). The phenotypes were not due to differences in electroporation or survival of DINs as GFP signal was similar between ATP1A3-WT or ATP1A3-E815K and they matching controls (Sup figure 4f and g). Altogether this further confirmed that modulating ATP1A3 activity levels impacts on DIN axon navigation and revealed pathogenic effect of the ATP1A3-E815K human mutation.

**Figure 6:**
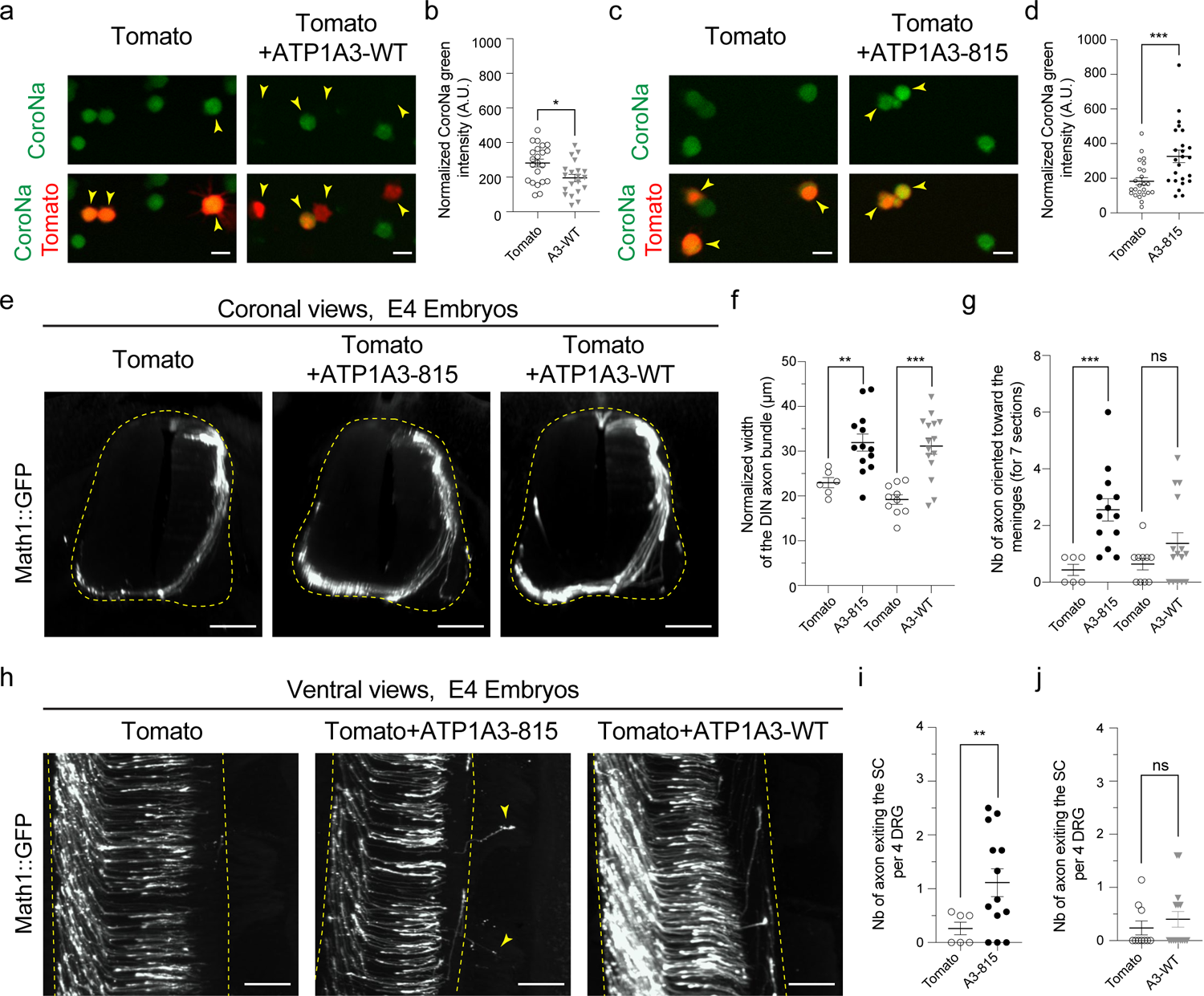
Overexpression of the E815K ATP1A3 form causing AHC disease impairs DIN axons navigation in the chicken spinal cord. (a) Microphotographs of the Na+ fluorescent indicator, CoroNa green, (green) and tomato (red) fluorescence in E3 dissociated electroporated DIN neurons. Images illustrate the decrease of CoroNa green fluorescence (lower intracellular Na^+^) in cell expressing tomato and ATP1A3-WT (arrowhead, right) compare to cells expressing tomato only (left arrowhead). (b) Scatter dot plot showing the normalized CoroNa green fluorescence intensity in tomato positive cells in tomato only and tomato+ ATP1A3-WT (A3-WT). Each point represents a field of view. (c) Microphotograph of CoroNa green fluorescence in tomato positive cells (arrowhead) in cells tranfected with plasmids expressing tomato alone (left) or with the ATP1A3 bearing the E815K mutation (ATP1A3-815) (right). (d) Scatter dot plot of the CoroNa green fluorescence in cells expressing tomato alone or tomato and the ATP1A3-E815K (A3-815), showing the increase of intracellular Na^+^ in the A3-815 condition. (e) Coronal views of E4 embryos co-electroporated with Math1::GFP and tomato, ATP1A3-WT or ATP1A3-815 illustrating the alteration of DIN axon trajectories (white). (f-h) Scatter dot plots showing the DIN axon bundle width (f) and the number of axons abnormally orientating toward the meninges (g) in tomato alone versus tomato and ATP1A3-WT (A3-WT) and tomato versus tomato and ATP1A3-815 (A3-815) in E4 embryos. (h) Ventral views of spinal cord exemplifying the increase in spinal cord exit of DIN axons (white) expressing the ATP1A3 bearing the E815K mutation (middle panel, yellow arrowheads) compare to DIN axon expressing tomato alone or tomato and the ATP1A3-WT.(i,j) Scatter dot plots showing the number of axons exiting the spinal cord in each condition. ns non-significant, *p < 0.05; **p < 0.01; ***p < 0.001 (Mann-Whitney). Yellow dashed lines outline the spinal cord in e and g. Scale bars: 10µm (a and c), 100µm (e and h)

## Discussion

Our study reveals a significant role of ATP1A3 in spinal cord neuronal differentiation. We found that ATP1A3 is crucial for the precise axon navigation of dorsal spinal cord neurons, forming commissural circuits with brain structures. The most severe AHC-causing human ATP1A3 mutation disrupts this function, leading to altered axon pathfinding. ATP1A3 functions by mediating the response to an embryonic electric field generated by a trans-neural-epithelial potential, guiding axons within spinal cord tissue. These findings bring insights into the roles of channels and pumps at early stages of the formation of neuronal circuits and possible etiological causes for some related neurodevelopmental diseases.

First, our study shows that Na^+^/K^+^ ATPases contribute to circuit formation in the embryonic spinal cord. Our findings combining analyses of the embryonic spinal cord using scRNAseq dataset in human and of expression patterns in chick show a conserved switch of ATP1A3 expression during neuronal differentiation. In chick spinal neurons, the protein localizes in the soma, axons and growth cones, consistent with Na^+^/K^+^ pumps detection in growth cones of cultured DRG neurons ^69^ and growth cone particles from fetal rat brain ^70^. We also found that Na^+^/K^+^ pumps are active in chick spinal neurons, consistent with previous reports in cultured differentiating neurons ^18,71^. Experimental manipulations of dorsal spinal cord neurons (DINs) in the chick embryo model allowed us to dissect the contributions of ATP1A3 during early neuronal differentiation. First, our data suggest that ATP1A3 is dispensable for axon initiation and elongation. Likewise, *in vivo*, neither ATP1A3 knock-down nor overexpression of transport defective ATP1A3 mutant prevented DINs to form and elongate axons. *In vitro*, DIN axon extension was not impaired by ouabain application, consistent with studies using DRG neurons ^72^ and iPS cells defective for ATP1A3^73^. This contrasts with axon growth during regeneration, found to depend on ATP1A1 and ATP1A2 and possibly ATP1A3 ^72,74,75^. It might rely on differences between mature and immature neurons and/or on processes specific to regeneration such as membrane resealing of injured axons thought to require Na^+^/K^+^ pumps to restore Na+ levels ^76^. Second and in contrast, we found that altering ATP1A3 activity *in vivo* by loss and gain of function both resulted in abnormal trajectories of navigating DIN axons. This reveals that ATP1A3-containing Na^+^/K^+^ pumps play a role in the process of axon navigation and extends reported neuro-developmental contributions of Na^+^/K^+^ ATPases across species, ranging from control of ventricle development in zebrafish^77,78^, neuronal specification during neurulation in amphibians ^22,23^ to migration of cortical neurons in rodent and brain morphogenesis in children ^27,28,79^.

Second, our investigation of the mechanisms of action of ATP1A3 during axon navigation suggests that it enables DIN neurons to orient their axons in an embryonic spinal cord electric field (EF). *In vitro*, we observed by altering Na^+^/K^+^ pumps activity and ATP1A3 gene expression that both are required for DIN axons to perceive EFs and preferentially orient toward the cathode. Such role in axon guidance might be general. On the one hand, this is consistent with early work that already suggested contribution of endogenous EFs to axon navigation ^80^. Later on, various neuron subtypes from different species were found to orient in EFs *in vitro*. Interestingly, their axons exhibit specificities in sensitivity, orientation and mechanisms of response ^35,46–48,57,58,81–90^. More recently, evidence from cultured retina were brought, which supports that a local dorso-ventral EF within the developping retina contributes to intraretinal guidance of some RGCs and optic nerve formation ^89,91^. On the other hand, embryonic EFs have been described in several species, at different stages and in various tissues, from the initial primitive streak to the retina, the skin, the limb, the intestine and the brain ^36,49–52,92–96^. Moreover, similarities between early avian and rodent embryos argue for evolutionary conserved bio-electric patterns during embryonic morphogenesis ^94,95,97^. Likewise, our characterization of transneural tube potential (TNTP) in the chicken embryo central nervous system echoes previous reports in xenopus and axolotl embryos ^50,52,54^.

Consistently, we found that electric potential drops between the surface and the lumen of the neural tube, likely generating a medio-lateral EF gradient that probably varies along the dorso-ventral axis due to differences in tissue thickness and previously suggested composition of ion channels and pumps ^89,98^. Interestingly, both experimentally decreasing the TNTP and altering ATP1A3 function resulted in DIN axon medio-lateral spreading, even leading to spinal cord exit. Thus, the TNTP may provide medio-lateral spatial coordinates to navigating DIN axons. In addition, we also found with our ectopic transplantation paradigm that altering TNTP or ATP1A3 expression in delocalized DIN neurons prevented their axons to orient towards intact spinal cord tissue, as normally do the control conditions. This reveals that EFs can guide the axons over large scales, consistent with the view of EFs being vectors of both short and long-range topographic information and providing a spatial blueprint that orchestrates embryonic development ^33–37,49,92,97,99–101^. It also suggests that ATP1A3 has a cell-autonomous contribution to EF-mediated DIN axon navigation.

Third, our results suggest that ATP1A3-mediated axon navigation might be impaired in ATP1A3 pathologies. ATP1A3 mutations cause various neurodevelopmental diseases, from Rapid-Onset Dystonia-Parkinsonism (RDP) to neuropediatric syndromes with earlier onsets such as CAPOS (cerebellar ataxia, areflexia, pes cavus, optic atrophy, and sensorineural hearing loss) and alternating Hemiplegia of childhood (AHC), ^26,29,32,102^. By changing the resting membrane potential and neuronal activity, they are thought to affect the activity of neuronal circuits leading to atrophy of brain regions ^103–106^. Our findings suggest that ATP1A3 syndromes could be rooted in earlier alterations, affecting the navigating axonal projections. This hypothesis is not only supported by ATP1A3 early expression but interestingly also by recent clinical observations. Brain malformations were observed in neonate patients with “developmental and epileptic encephalopathy” (DEE), “Dystonia, Dysmorphism, Encephalopathy, MRI Abnormalities, and no Hemiplegia” (D-Demø) syndrome or Congenital Hydrocephalus^27–29,31,79^. This includes defects of corpus callosum, a major brain inter-hemispheric connection, in ATP1A3-related DEE and AHC patients ^27,107,108^. More generally, our findings echo those obtained for developmental channelopathies, affecting channels that contribute to axon navigation^27,109^. This is the case of NDMA receptors, whose mutations, as ATP1A3, can lead to polymicrogyria and have reported axon guidance functions ^110,111^. Our work also suggests that alterations of ATP1A3 functions may affect regions outside the brain. Interestingly in mouse models, loss of Math1 expressing spinal DIN neurons, the population that we found in the chick embryo to use ATP1A3 for axon navigation, leads to motor deficits that resemble those observed in ATP1A3 E815K knock-in mouse ^112,113^.

Finally, the mechanisms by which ATP1A3-containing Na^+^/K^+^ pumps direct growth cone steering during axon navigation remain to be determined. EFs were reported to mediate directional movement of transmembrane proteins and their polarization at the surface of migrating cells and growth cones ^35,48,114–117^. Interestingly, the lateral mobility of Na^+^/K^+^ pumps observed in developing axons ^69^ and of ATP1A3 in mature neurons ^21,118,119^ suggest that the pumps could be relocalized in presence of EFs, as also predicted by theoretical models^117^. During EF-oriented cell migration, Na^+^/K^+^ pumps were shown to relocate to protrusions and to ensure maintenance of their polarity through local membrane potential hyperpolarization ^38^. Membrane potential could regulate actin cytoskeleton dynamics and adhesion complexes turnover ^38,120,121^. This could also be the case in growth cone protrusions, whose dynamics is essential to axon reorientation including in response EFs and depend on actin cytoskeleton and adhesion complexes turnover ^85,122,123,123,124^. Even if our results suggest that Na^+^/K^+^ pump functions depend on their ion transport activity, they could contribute to growth cone turning in other ways. Polarized distribution of Na^+^/K^+^ pumps could shape focal calcium signaling ^125^ that can direct growth cone turning in response to EFs ^35,126^. Lastly, the proposed local hyperpolarization resulting from polarized ATP1A3 activity could also affect membrane electrostatics, possibly promoting the stabilization of positively charged proteins such as Cdc42 as suggested during EF-induced yeast polarization^127^.

It also remains open how EF and extracellular axon guidance molecules cooperate to guide DIN axons. Several molecular cues were identified to act at successive steps of their navigation, orienting them ventrally, driving their progression towards the floor plate, across the midline and their rostro-caudal turning after crossing^128–130^. Possibly, the two mechanisms could act in synergy to strengthen the robustness of the pathway selection process. Alternatively, they could act with different precision scales, for instance EF providing global information and guidance molecules more accurate positional information. Finally, both mechanisms could be integrated to generate unique landmarks, for example with EF that could regulate the spatial distribution of guidance receptors within the growth cones.

Whatever the case, our study reveals unexpected contribution of ATP1A3 during the navigation of axonal projections, that may apply to many neuron types both in the central and peripheral nervous system. It also paves the way for studies of alternative etiologies in the context of ATP1A3-related diseases and more globally in channelopathies with early developmental onset.

## Acknowledgments

We thank E Stoeckli (Germany) for the pMath1::GFP plasmid and D Beclin for the pCAGEN-IRES-Tomato plasmid. We thank F Amblard and J-L Bessereau for constructive discussions. This work was funded by the LabEx CORTEX and DEVWECAN of Université de Lyon, within the program "Investissements d’Avenir" (ANR-11-IDEX-0007) operated by the French National Research Agency (VC), the Association Française contre les Myopathies (AFM) (VC, JH), and the Fondation pour la Recherche Médicale (VC, SD).

## Author contributions

Conceptualization: VC, JF

Methodology: VC, JF, SD, SC, JCC, OP, GM, RF, FM, MA

Investigation: SD, SC, RG, JCC, JF, OP

Project administration: VC, JF

Funding acquisition: VC, SD, JH

Supervision: VC, JF

Visualization: SD, JF, SC, JCC, VC

Writing – original draft: SD, JF, VC

Writing – review & editing: SD, JF, VC, SC, FM, JCC, OP, JH

## Declaration of interests

The authors declare no competing interests.

## Data Availability

Data available on request from the authors.

## Supplemental information

Sup Figure 1 to 4

Sup Table 1: Excel file containing the information about the single cell dataset used and their processing, including the marker genes used.

## Methods

### Egg incubation and embryo staging

Fertilized eggs provided by Élevage du Grand Buisson (Chabanière, France) were incubated at 38.5°C in humidified incubators (MIR 154 Sanyo Electric Co., LTD.) and staged accordingly to Hamburger and Hamilton ^131^.

### ATP1A3 RT-PCR from chick embryo spinal cord

For RT-PCR analysis, total RNA was extracted from ventral or dorsal spinal cords dissected from HH18/HH19 (E3), HH25 (E4) or HH28 (E5) chick embryos using the NucleoSpin RNA XS kit (740902-10, Macherey-Nagel). One µg of RNA was used for the reverse-transcription using RevertAid H Minus First Strand cDNA Synthesis Kit (K1632, Thermo Fisher scientific). PCR was performed using the Platinum Green Hot Start PCR 2X MasterMix (Invitrogen, Thermo Fisher scientific) with the following forward GCCTACGGACAGATCGGGATG and reverse GTTGCGGCGGAGGATGAGT. Primer pairs were designed to encompass at least one intron. Intron-Exon junctions of chick transcripts were defined by sequence homology with the mouse ones.

### Chicken ATP1A3 cloning and point mutation

Chicken ATP1A3 coding sequence was inserted upstream of the IRES sequence of the pCAGEN-IRES-tomato (named tomato) kindly provided by D. Beclin from Marseille by in Fusion recombination. Wild type chick ATP1A3 was PCR amplified from the cDNAs generated from E3 dorsal spinal cord (see RT-PCR section). Amplification was done with CloneAmp HiFi (639298, Takara Bio). Primers were designed using the Takara website (https://www.takarabio.com/learning-centers/cloning/primer-design-and-other-tools) based on the ATP1A3 published sequence (NM_205475.1). ATP1A3 primer specificity was checked by primer blast (NCBI). Primers include EcoRI site (underlined) and sequence homologous to the plasmid are in bold.

F: **GCTTGATATC<u>GAATT</u>**<u>C</u>ATGGGGGACAAAGGGGAG

R: **GGAGAGGGGC<u>GAATT</u>**<u>C</u>TTAATAATACGTTTCCTTCTCCA

PCR fragment and EcoRI digested pCAGEN-IRES-tomato plasmid were gel purified. The fragment was then inserted in the pCAGEN-IRES-tomato with the In-Fusion® HD Cloning Kit (639649, Takara bio) according to manufacturer recommendations to generate pCAGEN-ATP1A3-WT-IRES-Tomato (named ATP1A3-WT). ATP1A3 was sequence verified and compared to the chicken reference sequence (NM_205475.1). Two substitutions leading to amino acid changes that exist in other species were consistently found.

The E815K human mutation was generated by PCR directed mutagenesis. Human and chicken protein sequences were aligned and the Glutamic acid corresponding to the E815 was identified (in bold in the amino acid sequence: D L G T D M V P A I S L A Y **E** A A E).

Mutagenesis primers were designed on the Takara website to introduce the needed substitution (mutated codon is in bold). F: GGCCTAC**AAA**GCAGCCGAGAGTGACATCATGAAGC R: GCTGC**TTT**GTAGGCCAGAGAGATGGCCG.

The fragment was PCR amplified on pCAGEN-ATP1A3-WT-IRES-Tomato using CloneAmp HiFi enzyme. PCR product was gel purified and assembled to generated the pCAGEN-ATP1A3-E815K-tomato (named ATP1A3-815) accordingly to In-Fusion® HD Cloning Kit guidelines.

### Processing of single cell public human data sets

The following datasets published by Rayon and collaborators were download from GEO reposit (GSM5236520 human_CS14_brachial_rep1, GSM5236521 human_CS14_thoracic_rep1, GSM5236522 human_CS17_brachial_rep1, GSM5236523 human_CS17_brachial_rep2, GSM5236524 human_CS17_thoracic_rep1, GSM5236525 human_CS17_thoracic_rep2)(https://www.ncbi.nlm.nih.gov/geo/query/acc.cgi?acc=GSM5236520). Datasets were analyzed in R studio (R 4.2.2) with Seurat (4.3.0). Datasets were first processed individually. Seurat object was trimmed to keep cells that expressed more than 200 genes and genes that are expressed in at least 10 cells. Next, cells with low or high counts and features as well as high mitochondrial gene percentages were removed (800<nFeature<5000, nCount<20000 and %mt<6 or 6.5, Additional file 1). A first clustering was done to remove non-spinal cord cells. Data were log normalized (NormalizeData), variable features identified (FindVariableFeatures, method = "vst", nfeatures = 2000), scaled and PCA dimension reduction was performed (RunPCA). Elbow plot and cluster tree were used to choose the dimensions for the FindNeighbors and the resolution for the FindClusters (Louvain algorithm) function respectively. Clusters were identified according to a list of known markers including the one used by Rayon and collaborators^132^ (additional file 1). Blood, mesodermal and floor plate cells as well as peripheral neuroblasts and neurons were removed. Scaling, PCA and clustering were run on the subset of spinal cord progenitor and neuron clusters. UMAP and subtype annotations were done to check data prior to integration. Subset data were scaled before identifying anchors (FindIntegrationAnchors, anchor.features = 2000, normalization.method = "LogNormalize", reduction = "cca" and default values for others parameters) and performing the integration with the IntegrateData function. Datasets from CS14 and CS17 were integrated independently and afterward together. Clustering was done as described previously, a contaminating NCC cluster was removed from the integrated data. Final clustering was performed using 30 dimensions and a resolution of 0.9 for CS14+CS17 and 0.8 for CS17. Neuronal differentiation was assayed with module score function for the following list ("NEFL","NEFM","MAP2","GAP43","STMN2","TUBB","MAPT"). Progenitor status was confirmed with the module score for the list ("TOP2A","PCNA","VIM","CDC20","AURKB","CDK1","HES5","HES1","MKI67","SOX2"). Volcano plot was generated with EnhancedVolcano (Blighe, K, S Rana, and M Lewis. 2018. “EnhancedVolcano: Publication-ready volcano plots with enhanced colouring and labeling.” https://github.com/kevinblighe/EnhancedVolcano.) after calculating gene expression differences between progenitor and neurons with FindMarkers with low thresholds (logfc.threshold = 0.01, min.pct = 0.1). Volcano plot colors were changed using ggplot2. Neuron and progenitor subtype annotations were done using dot plots of known markers (additional file 1) and Ucell score (AddModuleScore_UCell) based on Rayon’s knowledge matrix (additional file 1). Other illustrations were done using DimPlot, VlnPlot, DotPlot and FeaturePlot.

### *In ovo* electroporation

*In ovo* electroporation of HH14/HH15 chick embryos was performed as described previously ^133,134^. pHB9::EGFP (Hb9 promoter) [TJ#96] was a gift from T Jessell (Addgene plasmid # 16275 ; http://n2t.net/addgene:16275 ; RRID:Addgene_16275) ^135^. The Math1::GFP plasmid was a gift from E Stoeckli ^136^. Endofree plasmids or siRNA were used at the following concentration: 1 μg/μl pHB9::GFP; 2 μg/μl pMath1::GFP; 0.05 μg/μl tomato; 0.4 μg/μl pCAGEN-GFP ^133^; 1.25 µg/μl siATP1A3 (custom siRNA, sense: CUCGUUAACGAGCGCCUCA[dT][dT], antisense: UGAGGCGCUCGUUAACGAG[dT][dT], Sigma) or siCtl (SIC001, Sigma), 1.3 µg/µl pCAGEN-ATP1A3-WT-IRES-Tomato, 1.3 µg/µl pCAGEN-IRES-Tomato, 1.3 µg/µl pCAGEN-ATP1A3-E815K-IRES-Tomato. Plasmids were diluted in H_2_O and the solution was injected into the lumen of the neural tube using picopritzer III (Micro Control Instrument Ltd., UK). Electrodes (CUY611P7-4, Sonidel) were placed along the back of the embryo, at the lumbo-thoracic levels, and 3 pulses (25V, 50ms, 500ms interpulse) were delivered by CUY-21 generator (Sonidel). Electroporated embryos were then incubated at 38.5°C.

### *In ovo* graft of neurons and surgical opening of the spinal cord

Electroporated embryos were harvested the next day (HH20-21, E3.5) in cold HBSS. Ventral or dorsal spinal cords were isolated from donor chick embryos electroporated with HB9::GFP or Math1::GFP respectively, cut into explants and kept in cold Neurobasal (21103-49, Gibco). Explants were grafted in stage-matched recipient embryos. Extraembryonic membranes were opened and a cross-shaped wound was performed at the base of the right hindlimb using insect pin (FST, 26002-10). The explant was then quickly introduced into the wound with a round-ended glass tool. Opening of the spinal cord was performed prior to grafting by making a longitudinal incision of skin and the roof plate with fine forceps. Surgical opening of the spinal cord was done at E3. The incision was made with insect pin, from two to three somites rostrally to the hindlimb bud to one somite caudally.

### HRC RNA-FISH on wholemount embryos

Whole mount HCR RNA-FISH ^137^was performed as described in Andre et al. (manuscript in revision) with the custom probes designed by Molecular instrument. For ATP1A1 and ATP1A3 oligos were chosen in specific sequences and do not bind to the other ATP1A RNAs.

**Table 1:**
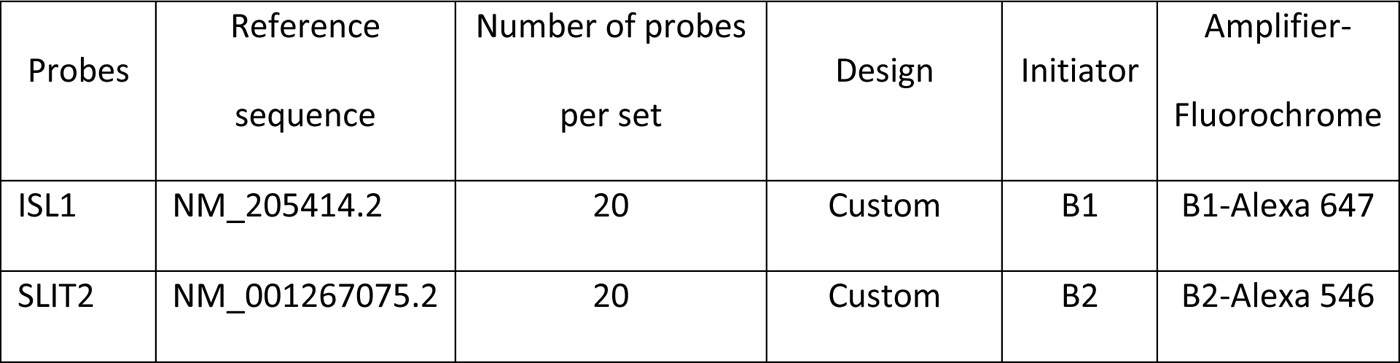
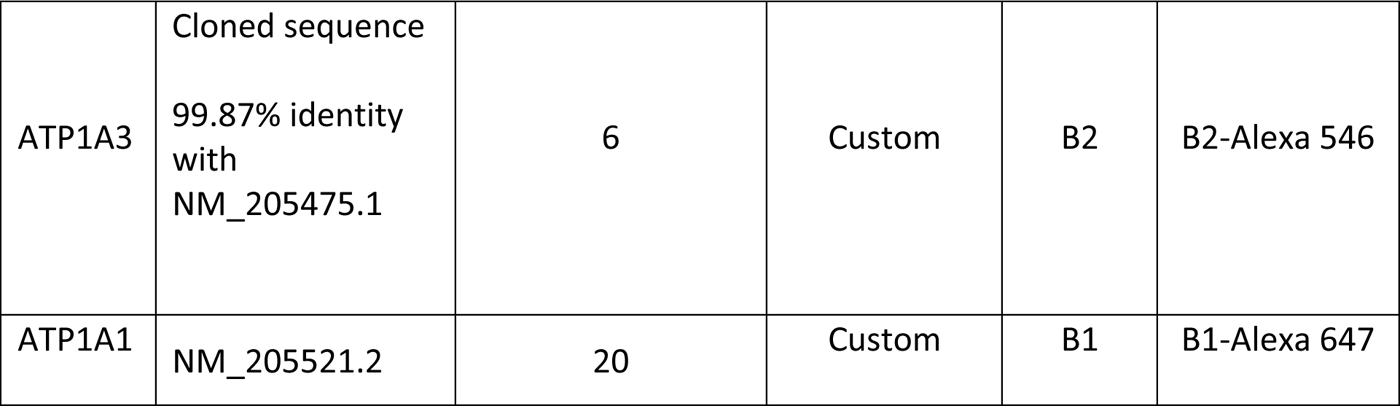
Probe list.

After dissection, embryos were fixed for 1 hour at room temperature (RT) in Paraformaldehyde (PFA) 4% in PBS 1X (32% stock, Electron Microscopy Sciences, 15714). They were washed 4 times for 10min in PBS 0.1% Tween20 (PBST) on ice before being dehydrated in MetOH/PBST baths (25%, 50%, 75%, 100%, 100% MetOH) for 30min on ice and were kept at -20°C. Embryos rehydrated in MetOH/PBST baths, (75%, 50%, 25%, 0%, 0% MetOH) and washed, twice in PBST, once in 50% PBST / 50% SSCT (Sodium Chloride Sodium Citrate 5X Tween20 0.1%) and once in SSCT. All washes were 5mins long and on ice. Samples were pre-hybridized in hybridization buffer (HCR^TM^ Buffers, Molecular instruments) for 30min at 37°C. Probe solutions were prepared at 4nM (e.g. 2pmol in 500µl in hybridization buffer) for ISL1, SLIT2 and ATP1A1 and 8nM for ATP1A3 and incubated with the embryos overnight at 37°C. Samples were washed 4 times for 15mins in the wash buffer (HCR^TM^ Buffers, Molecular instruments) at 37°C and twice in SSCT at RT. Hairpins were heated at 95°C for 90 seconds and cooled in the dark for 30min before being diluted in the amplification buffer (HCR^TM^ Buffers, Molecular instruments) at a final concentration of 30nM (15pmol for 500µl). Embryos were incubated with the amplification buffer for 5min at RT and overnight at RT with the amplification buffer containing the hairpins. The next day, samples were washed in SSCT at RT for 2 x 5min, 2 x 30min and 1 x 5min.

### Whole-Mount Immunofluorescence

Embryos were fixed in 4% PFA in PBS for 1h at room temperature before being permeabilized in the blocking solution (20% DMSO, 0.5% Triton X-100, 3% Bovine Serum Albumin and 100 mM glycine in PBS) for 24 hours. Next, the embryos were incubated with primary antibodies for four and half days. The embryos were washed with PBS containing 2% DMSO and 0.5% triton X100, 4 to 6 times over a day. Secondary antibodies diluted in the blocking solution were incubated overnight and samples were washed 4 to 6 times in PBS containing 2%DMSO and 0.5% triton for a day prior clearing. Primary and secondary antibodies were diluted at 1/500. The following primary antibodies were used: anti-GFP (A11122, Invitrogen, Thermo Fisher scientific) and anti-Neurofilament 160kDa (RMO-270, 130700, Invitrogen, Thermo Fisher scientific), RFP antibody (Rabbit IgG, AB62341, Abcam). Secondary antibodies used were: Goat anti Rabbit IgG Alexa 555 (A21429, Thermo Fisher scientific) and Donkey anti-mouse IgG FP 488 (FP-SA4110, Interchim), or Donkey anti-Rabbit IgG FP 647H (FP-SC5110, Interchim) and Donkey anti Mouse IgG Alexa 555 (A31570, Thermo Fisher scientific).

### Tissue Clearing and Selective Plane Illumination Microscopy (SPIM) Imaging

Tissues were cleared using modified Eci method ^138^. After immunolabeling, electroporated or grafted E4 and E5 embryos were washed in distilled water before being dehydrated by 30min incubations in 50%, 75% and 100% ethanol solutions. Additional 30 minutes and 2 hours incubations were made in absolute ethanol before placing the sample in Ethyl cinnamate (ECI,112372-100G, Sigma-Aldrich) overnight. All incubations were performed at room temperature in the dark on a roller mixer (roller mixer, Stuart equipment, SRT6D).

For HCR RNA FISH experiments sample were post-fixed in 4% PFA for 20min and washed 3 x 10min in PBS 1X at RT before being dehydrated on ice in MetOH/Ultrapure H_2_O baths, (20%, 40%, 60%, 80%, 100% MetOH) for 30min each. They were kept overnight in 100% MetOH before a last 3h 100% MetOH bath. Clearing was done in Ethyl Cinnamate (ECi). A first 1h ECi bath was done under soft agitation, followed by overnight ECi incubation.

Cleared samples were imaged using a light-sheet UltraMicroscope II and Blaze (LaVision Biotec, Miltenyi Biotec) equipped with macrozoom (0.063 to 6x) and 2x front objective or 4x front objective with 1x magnification lenses. When using the macro-Zoom the images were usually acquired at 3.2x magnification. Samples glued on the sample holder dorsal side up. Acquisition were performed with ImspectorPro (LaVision Biotec, Miltenyi Biotec) using ECI settings. Lightsheet thickness was set 4µm and images were acquired every 2µm. For some samples, Dynamic focus was used to kept Z-resolution constant over the region of interest.

### Image analysis, phenotype quantification and image preparation for illustration

Images series were converted in 3D images in Imaris 9.0.2 (Oxford instruments). Color, Intensity and contrast were adjusted using the display adjustment menu.

Axon trajectories from grafted DIN were analysed blindly in 3D view. Clipping planes and Orthoslicer were used to inspect the 3D volume in more details. This enables to count the axons in different regions and their orientation. Visual inspection was also used to attribute the guidance index value that represents the fraction of axons growing toward the limb over the one going toward the spinal cord.

DINs axon trajectories of E4 embryos (electroporated with 0Tomato, ATP1A3-WT or ATP1A3-815, Control siRNA or targeting ATP1A3) or E5 embryos (SiRNA and open spinal cord experiments) were analysed with Imaris 9.0.2 (Oxford instruments). In some cases, image stacks had to be rotated using the free rotate function to have consistent orientations for slice view. Number of axons extending outside the spinal cord were counted along the AP axis of the embryo and normalized for 4 ventral roots. To measure the DIN axon mediolateral distribution and orientation 6 to 8 extended sections of 10 µm (extended view in MIP mode) were generated regularly along the electroporated thoracic region: one at mid-DGR level and one in between DRG alternatively. Measurements of lengths and intensities were then done with FIJI software. Axon bundle width was measured and normalized to the spinal cord half width. GFP expression after electroporation was assessed by measuring the “Integrated density” of the DIN cluster in the dorsal spinal cord of the same sections. The number of axons getting toward the meninges were counted on the same sections and normalized with the mean of 7 sections. The images presented in the figures are snapshots of 3D view (MIP mode) or slice view (section with extended view in MIP mode).

### Primary culture of dissociated neurons

Embryos at HH19 to HH21 (E3-E3.5) were harvested in cold HBSS and the spinal cords were dissected. The collected ventral or dorsal spinal cords were trypsinized in HBSS containing 2.5 mg/mL of trypsin (T5266, Sigma-Aldrich) and 0.033mg/mL of DNAse1 (DN25, Sigma-Aldrich) at 37°C for 10min. Cells were mechanically dissociated using P1000 and P200.

Cells were plated on the glass coverslips for immunofluorescence (0.25×10^5^ cells), CellView glass bottom dishes (Cellview 627871, Greiner) for CoroNa green labeling (0.3×10^5^ cells) or in the Ibidi chambers (80106, µ-slide, Ibidi) for electric stimulation (0.6×10^5^ cells). Coverslips and chambers were coated with 50µg/ml of L-polylysine (P4707, Sigma-Aldrich) and 10µg/ml of laminin (L2020, Sigma-Aldrich). Dissociated neurons were cultured in Neurobasal (21103-49, Gibco) supplemented with 1% Penicillin/Streptomycin, 1X B27 (17504044, Gibco), 500µM glutamine (G7513, Sigma-Aldrich), 100ng/mL netrin (1109N1025, R&D Systems).

### Exposition to electric fields and analysis of axon orientation

The culture setup for EF exposure was similar to Pan and Borgens ^57^. Agarose 3% (A9414, Sigma-Aldrich) in Neurobasal was used to make bridges to connect the Ibidi chamber reservoirs to neurobasal-filled beakers containing the platinum electrodes (diameter 0.5mm, 267228, Sigma-Aldrich) connected to the power supply (EPS601, GE Healthcare/Amersham). Neurons were either exposed to EF during 24h starting 1 hour after plating, or during 8h after 16h of culture, without or with additional inhibitors. Ouabain (O3125, Sigma-Aldrich) was used at 10μM.

Cells were fixed with PBS supplemented with 4% paraformaldehyde, then permeabilized with 0.5% Triton 100X in PBS and incubated with 1µg/ml of phalloidin-TritC (P1151, Sigma-Aldrich) in PBS. Imaging was performed using an inverted microscope (Zeiss axiovert, 10X). The axon orientation was analyzed in blind, using ImageJ software by drawing straight lines from the nucleus to the growth cone. Only individual and clearly visible axons were used for analysis. The mean vector was calculated from the angle distribution of each culture to determine the mean angle and the strength of the response. Indeed, the norm of the vector or the orientation coefficient ranges from 0 (no response) to 1 (maximal response) ^139^.

### Intracellular Na^+^ and K^+^ measurements with fluorescent dyes

To test ouabain efficiency, neurons were treated 22h after plating with or without 10µM Ouabain (03125, Sigma-Aldrich) for 1 hour. Culture medium was then replaced by medium supplemented with 5µM CoroNa Green (CoroNa^TM^Green, C36676, Invitrogen, Thermo Fisher scientific) or 5µM CoroNa Green and 10µM Ouabain for 30 minutes. The cells were washed with Neurobasal and left either in Neurobasal or in Neurobasal with 10µM Ouabain for immediate live imaging.

Dorsal spinal cord cells from embryos electroporated with tomato, ATP1A3-WT and ATP1A3-815 plasmids were seeded on CellView dishes (627871,) at 3.10^5^ per compartments and left one hour at 37°C in Neurobasal and HBSS before adding respectively 5µM CoroNa Green or 10µM PBSI (P-1267, Molecular probes) previously mixed with Pluronic F-127 (final concentration 0.06%, P3000MP, Invitrogen). Cells were incubated for 30 minutes with the dyes and were rapidly washed with Neurobasal or HBSS respectively and imaging immediately. Imaging was performed at 37°c on Olympus IX81 microscope equipped with a spinning disk (CSU-X1 5000rpm, Yokogawa) and an Okolab environmental chamber. Images were acquired with a 20X objective (0.75 N.A) on EMCCD camera (iXon3 DU-885, Andor technology). Corona Green and PBSI were excited with 488nm laser and its signal collected using GFP filter (shamrock single band, 525nm centered, 50nm range). 4-compartments dishes were use so that control and Oubain treated cells as well as tomato and ATP1A3-WT or ATP1A3-815 expressing cell could be imaged simultaneously. Multi-Position Z-Stacks for the different wavelengths (DIC and GFP or RFP and GFP) were acquired with the “multidimensional acquisition” function of Metamorph. The images were analyzed with FIJI software. DIC or tomato channel was used to identify cells and to outline their soma with oval selection. CoroNa Green and PBSI signal intensity was measured within the outline and background intensity measured in regions without cell was subtracted.

### ATP1A3 labeling and SiRNA knockdown efficiency

Immunofluorescence were performed on PFA-fixed cultures (2% PFA, 20 minutes at room temperature). Dissociated cells were blocked in 6% BSA and immunolabeled with anti-α3 antibody (mouse Anti-ATP1A3 xVIF 9-G10, MA3-915, Invitrogen, Thermo Fisher scientific) for 2 hours 200 times in PBS1x containing 1%BSA or at room temperature. Cells were washed 3 times 10 min in PBS before being incubated with the appropriate anti-mouse secondary antibody (1:400 in PBS containing 3%BSA) for 1 hour at room temperature. ATP1A3 distribution in differentiated DINs was analyzed by confocal microscopy (40X, Olympus FV1000). ATP1A3 signal intensity in GFP and SiRNA transfected neuron was quantified from stack of confocal images (20X, Olympus FV1000) one hours and half after plating. ATP1A3 signal intensities were summed and quantified in the surface of the cell that was segmented after automatic threshold of the GFP signal. Quantifications were done in ImageJ.

### Immunofluorescent labelling on Cryosection

PFA-fixed embryos were embedded in gelatin/sucrose as previously described ^140^. 50-60 µm cryosections were blocked for 3h at room temperature in PBS containing 3% BSA (A7906, Sigma-Aldrich) and 0.5%Triton X-100 (T9284, Sigma-Aldrich). Slices were incubated for 48h at 4°C with primary antibodies anti-GFP (A11122, Invitrogen, Thermo Fisher scientific, 1:500), anti-Robo3 human (AF3076, R&D, 1:100), anti-neurofilament-160kDa (RMO270, 130700, Thermo Fisher scientific, 1:500), anti-Islet1/2 (39.4D5, DSHB, 1:500) anti-GFP (Ab5450, Abcam, 1:500), diluted in the blocking solution. Appropriated secondary antibodies anti-rabbit (A31573, MolecularProbes), anti-mouse (50185, Sigma-Aldrich; A21426, Thermo Fisher scientific) and anti-goat (A11055, Molecular Probes) diluted at 1:500 in PBS-3%BSA-0,5% triton were incubated 48h at 4°C. Slices were then washed, PFA-post-fixed and mounted in Mowiol. Staining was observed under Olympus FV1000 confocal microscope and images were analyzed with ImageJ.

### Electrophysiology

E4 embryo preparation was done similarly to Hotary and collaborators ^51^. After removal of 5ml of albumen, the egg shell was progressively removed to access to the embryo without damaging the area vasculosa. The extraembryonic blastoderm was cut outside the area vasculosa. The embryo with its area vasculosa was washed in PBS and then placed in center-well organ culture dish (353037, Falcon) whose well was filled with Sylgard (Sylgrad 184; Dow corning). PBS was gently removed to immobilized the embryo on top of the Sylgard. Only undamaged embryos were used, when needed extraembryonic membrane were removed and the dorsal roof plate of the hindbrain was opened with fine forceps and insect pins. After surgical incision the embryo were quickly washed 3 times with PBS. Recording were done just after embryo preparation at room temperature under a microscope (axiovert, Zeiss) to visually monitor the impalement progression. Tungsten monopolar electrodes (30021, Phymed) were used to measure the potential in rostral and caudal tectum and moved through the tissue with micromanipulators. Crocodile clip contacting the PBS left in the outside compartment of the dish was used as ground electrode. Electrodes were connected to AC/DC differential amplifier set at a gain of 50 (model 3000, A-M System^TM^) and the signal was filtered at 1 kHz low pass and 0.1 Hz high pass, measured potential differences were displayed and recorded using a homemade matlab program.

### Statistical analysis

Mann-Whitney, Wilcoxon matched-pairs and Chi² tests were performed using Prism 9.

## Notes

### Competing Interest Statement

The authors have declared no competing interest.

### Summary of Updates

Addition of the supplemental data file missing in the original automatic submission

