## supplemental figures for "Contribution of the neuron-specific ATP1A3 to embryonic spinal circuit emergence"

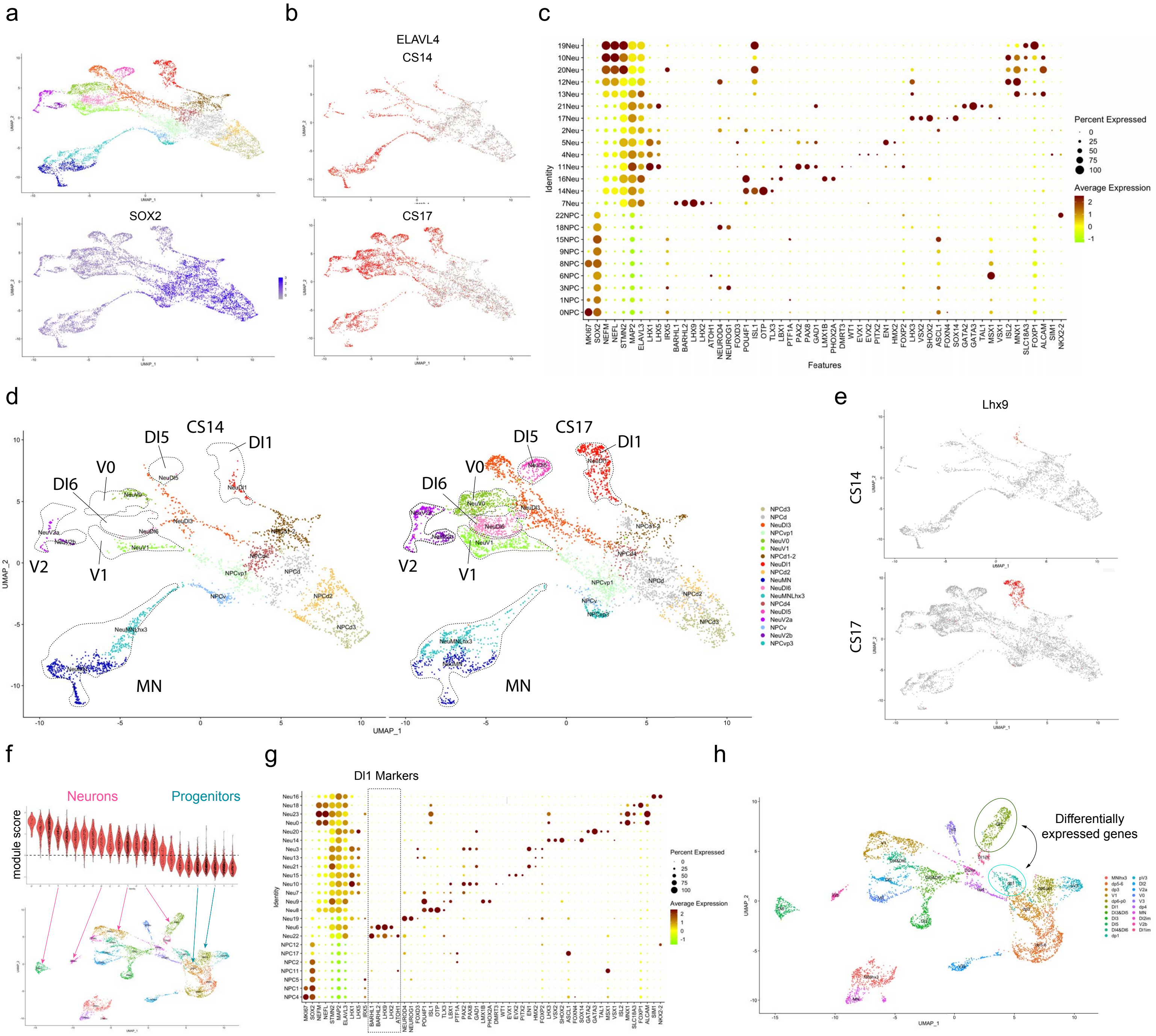
a
b
ELAVL4
CS14
SOX2
CS17
c
Percent Expressed
0
25
50
75
100
Average Expression
2
1
0
-1
Identity
Features
19Neu
10Neu
20Neu
12Neu
13Neu
21Neu
17Neu
2Neu
5Neu
4Neu
11Neu
16Neu
14Neu
7Neu
22NPC
18NPC
15NPC
9NPC
8NPC
6NPC
3NPC
1NPC
0NPC
MKI67
SOX2
NEFM
NEFL
STMN2
MAP2
ELAVL3
LHX1
LHX5
IRX5
BARHL1
BARHL2
LHX9
LHX2
ATOH1
NEUROD4
NEUROG1
FOXD3
POU4F1
ISL1
OTP
TLX3
LBX1
PTF1A
PAX2
PAX8
GAD1
LMX1B
PHOX2A
DMRT3
WT1
EVX1
EVX2
PITX2
EN1
HMX2
FOXP2
LHX3
VSX2
SHOX2
ASCL1
SOX14
GATA2
GATA3
TAL1
MSX1
VSX1
ISL2
MNX1
SLC18A3
FOXP1
ALCAM
SIM1
NKX2-2
d
CS14
CS17
DI5
DI1
V0
DI6
V2
V1
MN
UMAP_1
UMAP_2
NPCd3
NPCd
NeuDI3
NPCvp1
NeuV0
NeuV1
NPCd1-2
NeuDI1
NPCd2
NeuMN
NeuDI6
NeuMNLhx3
NPCd4
NeuDI5
NeuV2a
NPCv
NeuV2b
NPCvp3
e
Lhx9
CS14
CS17
f
Neurons
Progenitors
module score
g
DI1 Markers
h
Differentially expressed genes
UMAP_1
UMAP_2

Sup Fig. 1

**Sup Figure 1: Clustering of single cell RNAseq of human spinal cord at CS14 and CS17 and characterization of neuron subtypes.**

(a) UMAP showing the different clusters identified by unsupervised SEURAT clustering after integration of CS14 and CS17 samples (top) and Feature plot illustrating the expression of the progenitor marker SOX2. (b) Feature plots showing the expression of the neural marker ELAVL4 at CS14 and CS17 to illustrate the increase of neuronal specification between the two stages. (c) Dot plots presenting the relative expression of known marker genes for progenitor and neuron subtypes. Dot color indicate the relative expression level and its size the percentage of cells with non-zero expression among the cluster. (d) UMAP of the CS14 and CS17 data showing the cluster identities defined with the markers presented in c and illustrating the appearance of several neuron clusters at CS17 (Dl5 and Dl6). (e) Feature plots showing the increase of cells expressing the DI1 marker LHX9 between CS14 and CS17 (f) Violin plots of the module scores for the neuronal signature in the different clusters identified at CS17 (top). Cluster with average expression above the dashed line (0 score) were categorized as neuronal. The two clusters with a module score close to 0 were classified as immature neurons. Arrows exemplify the neuronal and progenitor annotations of some clusters shown on the UMAP of CS17 cells. (g) Dot plot of the relative expression of known marker genes for progenitor and neuron subtypes in CS17 clusters. DI1 markers are highlighted with the dashed line box (h) UMAP annotated with the subtype identities defined based on the marker expressions shown in g. Circles indicate the DI1 neuron and the DI1-2 dorsal progenitor cluster used to quantify the increase of ATP1A3 expression in Dl1 neurons.

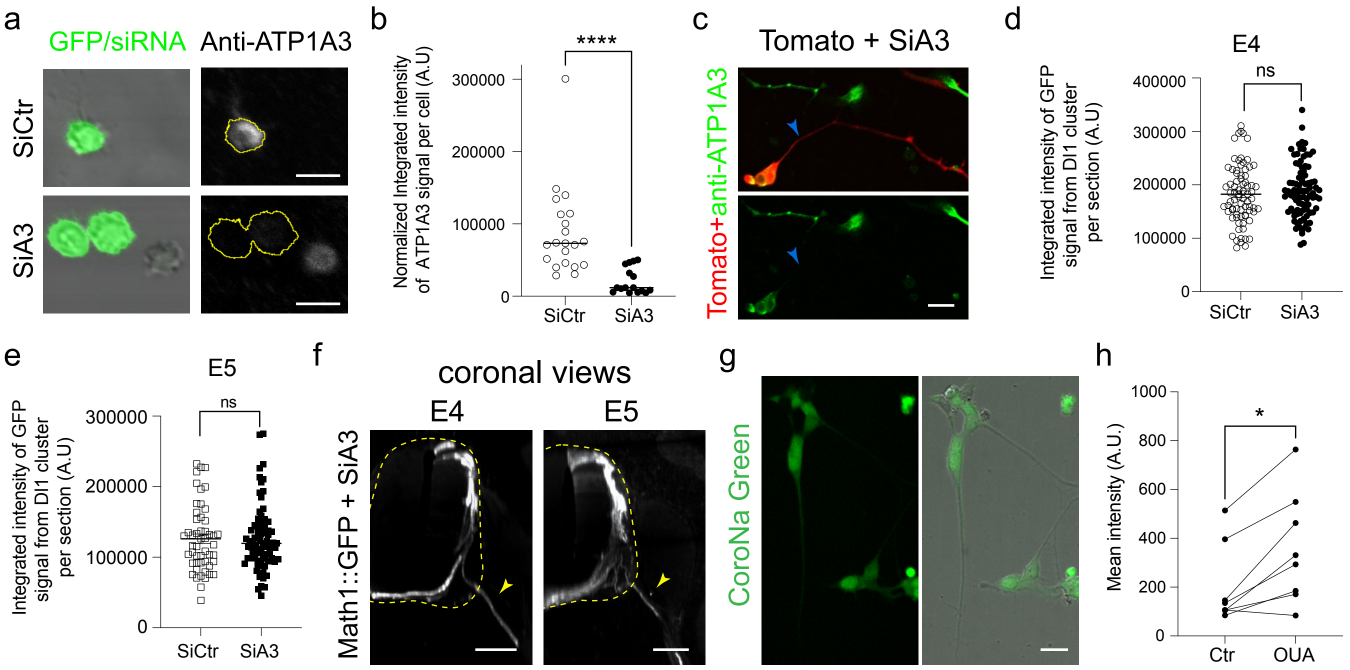
a
GFP/siRNA
Anti-ATP1A3
SiCtr
SiA3
b
Normalized Integrated intensity of ATP1A3 signal per cell (A.U)
****
300000
200000
100000
0
SiCtr
SiA3
c
Tomato + SiA3
Tomato+anti-ATP1A3
d
E4
Integrated intensity of GFP signal from DI1 cluster per section (A.U)
ns
400000
300000
200000
100000
0
SiCtr
SiA3
e
E5
Integrated intensity of GFP signal from DI1 cluster per section (A.U)
ns
300000
200000
100000
0
SiCtr
SiA3
f
coronal views
E4
E5
Math1::GFP + SiA3
g
CoroNa Green
h
Mean intensity (A.U.)
*
1000
800
600
400
200
0
Ctr
OUA

Sup Figure 2

**Sup Figure 2: SiRNA electroporation decreases ATP1A3 expression without affecting Math1::GFP expression and Ouabain inhibit $Na^+$ transport.**

(a) Images of the α3 immunolabeling on cells electroporated with GFP encoding plasmid and either control siRNA (SiCtr) or the siRNA targeting ATP1A3 (SiA3), 1h30 after plating. The left panels show the phase and GFP signal. The right panels show the α3 immunolabeling whose intensity was measured in the yellow cell outline defined from the GFP signal. (g) Scatter dot plot showing fluorescence intensity of α3 immunolabeling in dissociated neurons electroporated with scrambled siRNA (Ctr) or siRNA targeting ATP1A3 (SiA3). (c) anti-ATP1A3 immunolabeling of dissociated neurons electroporated with siRNA against ATP1A3 (SiA3) and tomato coding plasmid after 1 day in vitro. Arrows point to the axon of a tomato+ neuron exhibiting markedly decreased anti-ATP1A3 staining compared to non-transfected cells. (d-e) Scatter dot plots showing the level of GFP expression in the DI1 cluster electroporated with SiCtr and SiA3 in E4 embryos (d) and E5 embryos (e). (f) Coronal views of E4 (left) and E5 (right) spinal cord showing DIN axons (white) that had exited the spinal cord (arrowheads). Dashed lines outline the spinal cord. (g) Microphotographs of the $Na^+$ fluorescent indicator, CoroNa green staining (green) in dissociated DIN neurons. The right panel shows the overlay of the CoroNa green signal and the DIC image. (h) Scatter dot plot showing the mean normalized intensity of the CoroNa green fluorescence for cultures treated with Ouabain (OUA) or not (Ctr). Lines link Ouabain-treated cultures to their matching control culture that were imaged together. (Ctr: 198.9 ± 57.42, N=8 experiments; OUA: 354.9 ± 79.89, N=8 experiments, p=0.0156). ns non-significant, *p<0.05, ****p < 0.0001; (Mann-Whitney in b, d and e Wilcoxon matched-pairs in h). Scale bars: 10µm (a), 20µm (c and g), 100µm (f).

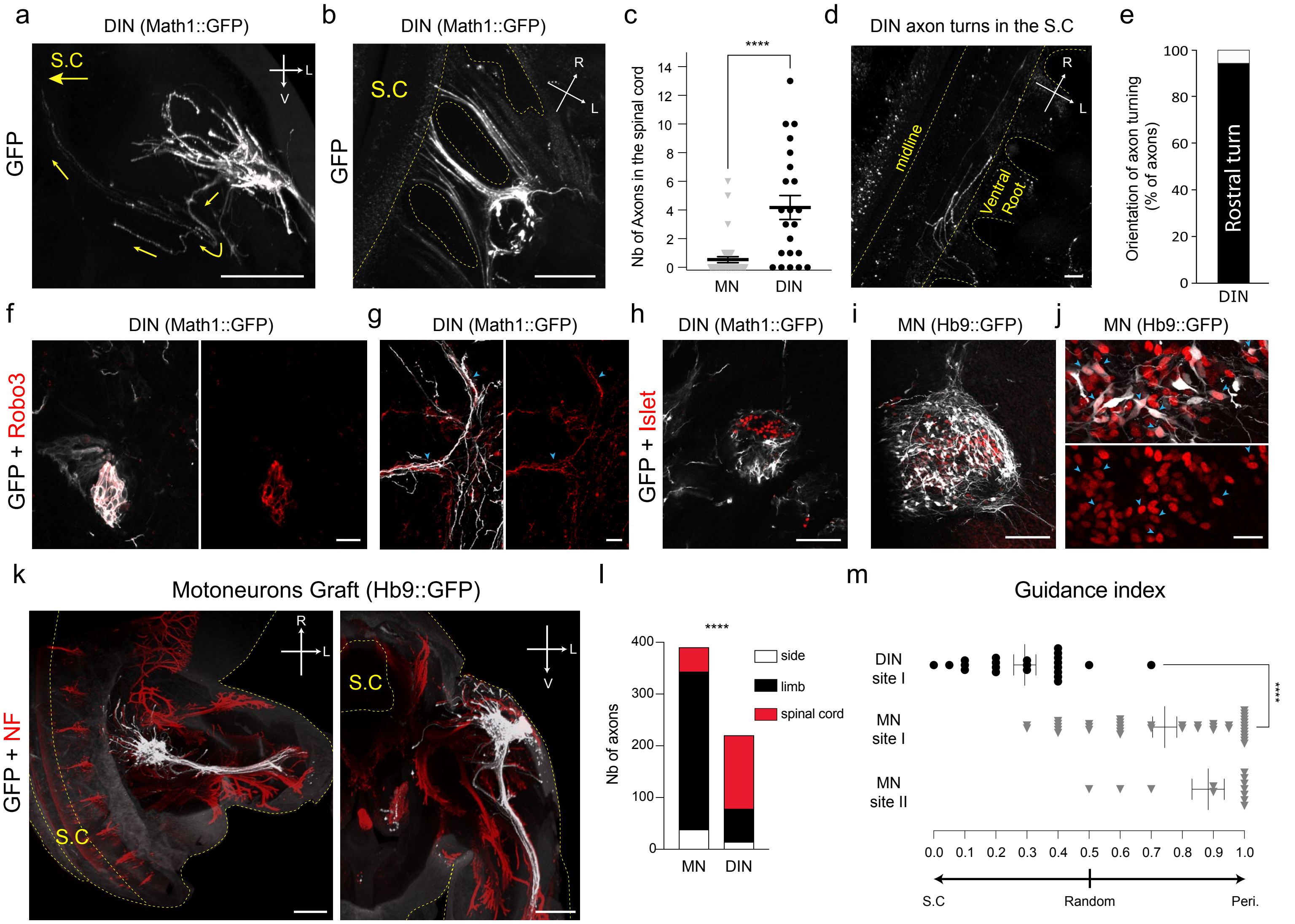
a
DIN (Math1::GFP)
GFP
S.C
b
DIN (Math1::GFP)
GFP
S.C
c
Nb of Axons in the spinal cord
****
MN
DIN
d
DIN axon turns in the S.C
midline
Ventral Root
e
Orientation of axon turning (% of axons)
Rostral turn
DIN
f
DIN (Math1::GFP)
GFP + Robo3
g
DIN (Math1::GFP)
h
DIN (Math1::GFP)
GFP + Islet
i
MN (Hb9::GFP)
j
MN (Hb9::GFP)
k
Motoneurons Graft (Hb9::GFP)
GFP + NF
S.C
l
Nb of axons
****
side
limb
spinal cord
MN
DIN
m
Guidance index
DIN site I
MN site I
MN site II
****
S.C
Random
Peri.

Sup Figure 3

**Sup Figure 3: Grafted DINs retain their identity and their axons exhibit DIN specific behaviors.**

(a) Coronal view of GFP positive DIN axons (white) changing their trajectories to orient toward the spinal cord. Arrows indicate axon orientations and spinal cord direction (top, S.C). (b) Images of DIN axons (white) entering the spinal cord by the ventral roots. (c) Scatter dot plot of the number of MN (grey triangles) and DIN axons (black dots) entering the spinal cord/graft. Lines indicate mean and SEM. (d) Images of DIN axons illustrating their preferential rostral turn in the spinal cord. Dashed lines outline the ventral roots, the spinal cord (S.C) boundaries (b and d) and the midline (d). (e) Histogram of the percentage of DIN axon turning in rostral (n=48; 94.3%) or caudal (n=3; 5.7%) direction. (f and g) Microphotographs exemplifying the co-expression of Robo3 (red) and GFP (white) within a dorsal spinal cord explants two days after grafting (f) and in some of the GFP+ axons formed by grafted neurons (g) (blue arrowheads). (h) Microphotograph showing the segregation of the DI3 marker, Islet 1/2 (red) and GFP (white) domains in grafted dorsal explant electroporated with the Math1:: GFP (DI1). (i and j) Microphotographs of ventral spinal cord explant two days after grafting showing the co-expression of the GFP (white) under the control of HB9 and Islet 1/2 (red) that is also expressed in motoneurons (MN) within the same region of the explant (i) and the same cells (j) (blue arrowheads). (k) Top and coronal views of grafted MNs (HB9::GFP) in cleared whole-mount embryos exemplifying MN axons preference for the limb. GFP staining is shown in white and neurofilament (NF) in red. Yellow dashed lines outline the embryo and the spinal cord (S.C). (l) Cumulative histograms of the number of axons orientating towards the limb, the spinal cord or along the rostro-caudal axis (side) (MN grafts (N=39): 38 side – 305 limb – 47 S.C; DIN grafts (N=22): 14 side– 64 limb – 142 S.C; Chi-square test). (m) Scatter dot plots showing the guidance index of axons formed by MN (grey triangles) and DIN explants (black dots) after grafting. Lines indicate mean and SEM. ****$p < 0.0001$ (Mann-Whitney in b and m, Chi-square test in l). R, rostral; L, lateral, V ventral, S.C Spinal cord. Scale bars: 300µm (a and k), 100µm (b, d, h and i), 20µm (f, g and j).

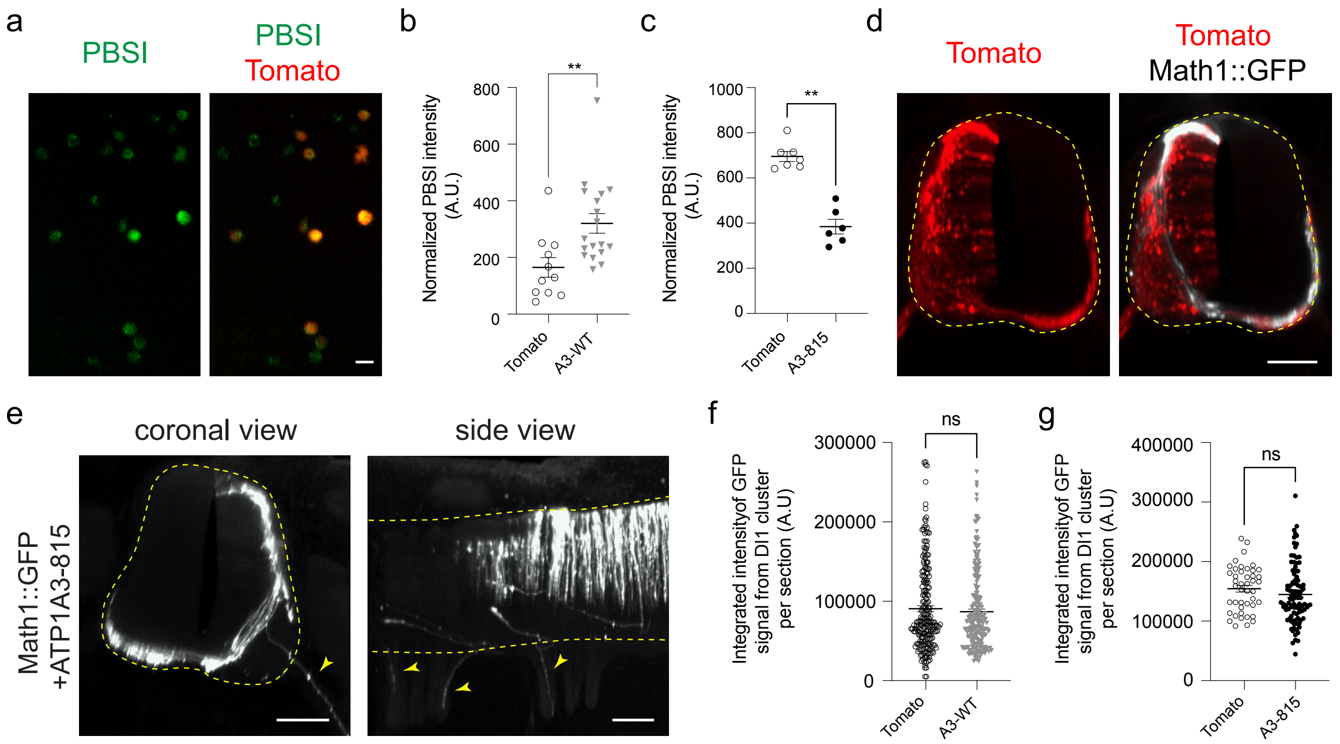
a
PBSI
PBSI
Tomato
b
Normalized PBSI intensity (A.U.)
800
600
400
200
0
**
Tomato
A3-WT
c
Normalized PBSI intensity (A.U.)
1000
800
600
400
200
0
**
Tomato
A3-815
d
Tomato
Tomato
Math1::GFP
e
coronal view
side view
Math1::GFP +ATP1A3-815
f
Integrated intensityof GFP signal from DI1 cluster per section (A.U)
300000
200000
100000
0
ns
Tomato
A3-WT
g
Integrated intensityof GFP signal from DI1 cluster per section (A.U)
400000
300000
200000
100000
0
ns
Tomato
A3-815

Sup Figure 4

**Sup Figure 4: ATP1A3-WT and ATP1A3-E815K expression effects on intracellular K+ concentration, Math1::GFP expression and spinal cord exit of DIN axons.**

(a) Microphotographs of the $K^+$ fluorescent indicator, PBSI (green) and tomato (red) fluorescence in live E3 dissociated DINs. The right panel shows the merge. (b-c) Scatter dot plots showing the PBSI intensity per field for the different conditions: Tomato with ATP1A3-WT (A3-WT) (b) or ATP1A3-E815K (A3-815) (c) and their respective matching control (Tomato alone) (d) Coronal views of the spinal cord showing the expression of tomato (red) and GFP (white) after Math1::GFP electroporation. The right panel shows the merge and dashed lines outline the spinal cord. (e) Coronal and side view of E4 embryos electroporated with Math1::GFP and ATP1A3-815. Yellow arrows show the axons exiting the spinal cord and dashed lines outline the spinal cord. (f-g) Scatter dot plots showing the GFP signal quantified on coronal sections for tomato, ATP1A3-WT and ATP1A3-815 electroporated embryos at E4. Lines on scatter dot plots indicate mean and SEM. ns non-significant, **$p < 0.01$ (Mann-Whitney). Scale bars: 10µm (a), 100µm (d and e).
